# Metabolite exchange within the microbiome produces compounds that influence *Drosophila* behavior

**DOI:** 10.1101/066035

**Authors:** Caleb N. Fischer, Eric Trautman, Jason M. Crawford, Eric V. Stabb, Nichole A. Broderick, Jo Handelsman

## Abstract

Animals host multi-species microbial communities (microbiomes) whose properties may result from inter-species interactions; however current understanding of host-microbiome interactions is derived mostly from studies in which is it is difficult to elucidate microbe-microbe interactions. In exploring how *Drosophila melanogaster* acquires its microbiome, we found that a microbial community influences *Drosophila* olfactory and egg-laying behaviors differently than individual members. *Drosophila* prefers a *Saccharomyces*-*Acetobacter* co-culture to the same microorganisms grown individually and then mixed, a response mainly due to the conserved olfactory receptor, *Or42b*. *Acetobacter* metabolism of *Saccharomyces*-derived ethanol was necessary, and acetate and its metabolic derivatives were sufficient, for co-culture preference. Preference correlated with three emergent co-culture properties: ethanol catabolism, a distinct volatile emission profile, and yeast population decline. We describe a molecular mechanism by which a microbial community affects animal behavior. Our results support a model whereby emergent metabolites signal *Drosophila* to acquire its preferred multispecies microbiome.

## Introduction

Multispecies microbial communities (microbiomes) influence animal biology in diverse ways (1): microbiomes modulate disease (2), metabolize nutrients (3), synthesize vitamins (4), and modify behavior (5). A central goal in host-microbiome studies is to understand the molecular mechanisms underpinning these diverse microbiome functions.

Some aspects of microbial community function are the product of interspecies interactions (6-9). For example, microorganisms modulate the metabolomes of neighboring species (10) and microbial metabolites (e.g., antibiotics) alter bacterial transcriptional responses (11). Despite current understanding of microbial inter-species interactions *in vitro*, some of which has been elucidated in exquisite detail, the consequences of microbial interspecies interactions within host-associated microbiomes are just beginning to be explored experimentally.

Insight into host-associated microbiome function has stemmed mostly from whole-microbiome (e.g., re-association of germ-free hosts with whole microbiomes (12)) and modeling microbiome function based on gene annotation (13) or single-microorganism (e.g., re-association of germ-free hosts with a single microorganism (14)) studies. However, these approaches tend to reveal only limited insight into interspecies microbial interactions, which can provide hosts with essential services. For example, termite symbionts carry genes necessary for metabolism of different parts of complex carbohydrates (15), yet their function has not been demonstrated *in vivo*; co-occurring human gut symbionts share polysaccharide breakdown products cooperatively (16, 17), but the consequences of such interactions for the host are unknown; inter-species bacterial interactions protect *Hydra* from fungal infection (18), but the mechanism of host protection is unclear. The need to understand the effects of inter-species microbiome interactions motivated our current work.

Attractive model systems in which to study the outcomes of inter-species microbial interactions for host biology would include a tractable host that harbors a simple multispecies microbiome. Here, we report the use of *Drosophila melanogaster* to study interactions in a simple microbiome and their consequences for host behavior.

The *Drosophila* microbiome consists largely of yeasts, acetic acid bacteria, and lactic acid bacteria (19-22). *Drosophila* ingests microbiome members from the environment (e.g., fermenting fruit, (22-26)), a behavior posited as a mechanism for *Drosophila* to select, acquire, and maintain its microbiome (21, 25). *Drosophila* behavior toward environmental microorganisms has focused on yeasts (27-31). While yeasts attract *Drosophila* via ester production (28, 29), lactic and acetic acid bacteria produce metabolites (e.g., acids) that may repel *Drosophila*, based on olfactory avoidance of high acid concentrations (32). One motivation of our study was to analyze *Drosophila* behavior toward the yeast and bacteria that dominate the *Drosophila* microbiome.

Yeast and bacteria are largely studied within separate *Drosophila* sub-disciplines, despite their shared habitat (21). Yeasts serve as food, providing *Drosophila* vitamins, sterols, and amino acids (21). Lactic and acetic acid bacteria are gut microbiome members (33) promoting larval development (34, 35), increasing resistance to pathogens (25), inducing intestinal stem cell proliferation (36), and reducing adult sugar and lipid levels (37, 38). Since microorganisms that are traditionally considered ‘food’ co-exist with those considered “microbiome’ in fruit fermentations and the two groups provide *Drosophila* with different services, we hypothesized that *Drosophila* might detect a beneficial community via metabolites that are produced cooperatively by the desirable symbionts. Alternatively, *Drosophila* might detect a different metabolite as the signal for each symbiont.

Fruit undergoes a well-characterized ripening process in which cell-wall degrading enzymes and amylases convert the firm, starchy tissue into soft, sugar-rich fruit (39-41). The high sugar content supports microbial colonization and fermentation by *Drosophila*-associated microorganisms, including yeasts, lactic acid bacteria, and acetic acid bacteria (23, 42). *Drosophila* avoids ‘green’ fruit and is attracted to ‘overripe’ fruit (43), yet it is unclear whether *Drosophila* behavior is influenced by the fruit microbiome and its metabolic properties. To this end, we developed a model fruit fermentation system that afforded measurement of microbial populations, microbial metabolites, and *Drosophila* behavior.

Here we demonstrate the importance of emergent microbiome metabolism—quantitatively different or unique metabolites produced by the microbiome, but not by any of its members in isolation—on behavior, suggesting that *Drosophila* modulates its chemical intake by selecting a beneficial microbiome.

## Results

To determine whether *Drosophila* responds to emergent microbial community metabolites, we used the T-maze olfactory assay to analyze *Drosophila* behavioral responses to several *Drosophila* microbiome members grown individually or in communities (Figure 1A, Table S1, Figure 1-figure supplement 1). When the strains were grown individually, *Drosophila* was strongly attracted to yeasts, moderately attracted to acetic acid bacteria, and neutral or slightly repelled by lactic acid bacteria (Figure 1B, C). Because strains within a microbial group attracted *Drosophila* similarly, a representative yeast, acetic acid bacterium, and lactic acid bacterium were used to test the effect of interactions between microbiome members on *Drosophila* behavior. *Drosophila* preferred microbial communities grown together to microorganisms grown individually and then mixed prior to analysis (defined throughout as a separate-culture mixture, Figure 1D). Focusing on a model *Saccharomyces cerevisiae* and *Acetobacter malorum* community, we found that when tested against apple juice medium (AJM), *Drosophila* preferred the community to the separate-culture mixture or individual members (Figure 1E). In sum, *Drosophila* detects, and prefers, microorganisms growing together to a mixture of the same strains combined after they had completed growth.

**Fig. 1.**
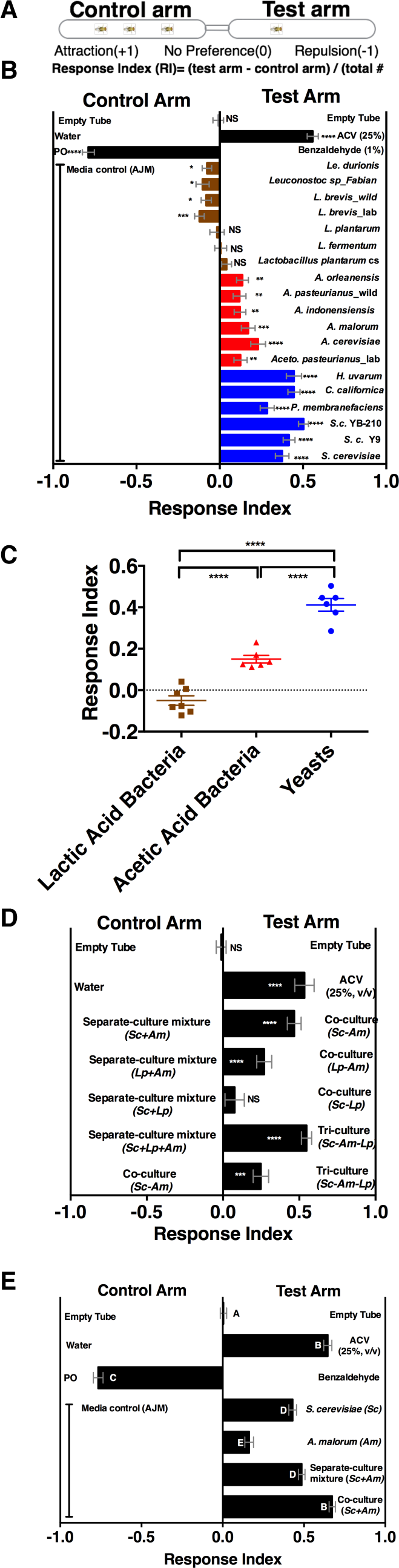
*Drosophila* detection of microbe-microbe metabolic exchange. **(A)** T-maze setup and quantification. **(B)** *Drosophila* behavior toward yeasts (blue), acetic acid bacteria (red), and lactic acid bacteria (brown) (Table S1). Mean ± SEM of 12-36 replicates (n=2-6 experiments). Each T-maze replicate uses a technical replicate of a microbial culture and one cohort of *Drosophila* maintained in separate vials for 3-5 days. Mock (two empty tubes), ACV (25% apple cider vinegar versus water), and benzaldehyde (1% versus paraffin oil (PO)). A one-sample t-test assessed deviance from 0. Symbols: NS p >0.05; * p ≤0.05; ** p ≤0.01; *** p ≤0.001; **** p ≤0.0001. (**C)** Mean *Drosophila* behavior toward each microorganisms was graphed according to microbial group and analyzed by one-way ANOVA. **(D)** *Drosophila* behavior toward community combinations of a representative yeast, acetic acid bacterium, and lactic acid bacterium in relation to their separate-culture mixture (grown individually and mixed; *Sc = S. cerevisiae*; *Am= A. malorum; Lp = L. plantarum* cs) grown for 96 h; *Drosophila* preference for the three-versus two-membered community is the last column. Mean ± SEM of 12-18 replicates (n=2-3 experiments). A one-sample t-test assessed deviance from 0. **(E)** *Drosophila* olfactory behavior toward the *S. cerevisiae* and *A. malorum* community and its constituent parts relative to media grown for 48-60 h. Mean ± SEM of 18-30 replicates (n=5 experiments). Differences between groups were assessed by ANOVA. A Tukey’s multiple comparison test was used to assess significant differences between groups (p≤0.05), as indicated by unique letters.

We next measured the attractiveness and other properties of the co-culture over time. When grown alone, the microorganisms had similar growth profiles (Figure 2A). However, when grown with *A. malorum, S. cerevisiae* populations first increased, then decreased between 60 and 72 h, and were undetectable by 96 h (Figure 2A). The decrease in *S. cerevisiae* viable counts mirrored a decrease in pH (Figure 2-figure supplement 1A). *Drosophila* did not prefer the co-culture relative to the separate-culture mixture at 34 h, slightly preferred the co-culture at 48 and 54 h, and strongly preferred the co-culture at 60 through 96 h (Figure 2B). *Drosophila* preference for the co-culture correlated with lower pH and *S. cerevisiae* populations, despite *Drosophila* olfactory avoidance of acid (32) and reliance on yeast for nutrition (44) (Figure 2C, D). *Drosophila* preference did not correlate with viable *A. malorum* populations (Figure 2-figure supplement 1B). *Drosophila* preference for the co-culture increased relative to sterile media during 34-96 h of growth, which is consistent with the increase in *Drosophila* attraction being due to a property of the co-culture rather than to a decrease in attraction to the separate-culture mixture (Figure 2-figure supplement 1C). In sum, several properties of the microbial community (e.g. *S. cerevisiae* density, pH) parallel *Drosophila* detection of, and preference for, the co-culture.

**Fig. 2.**
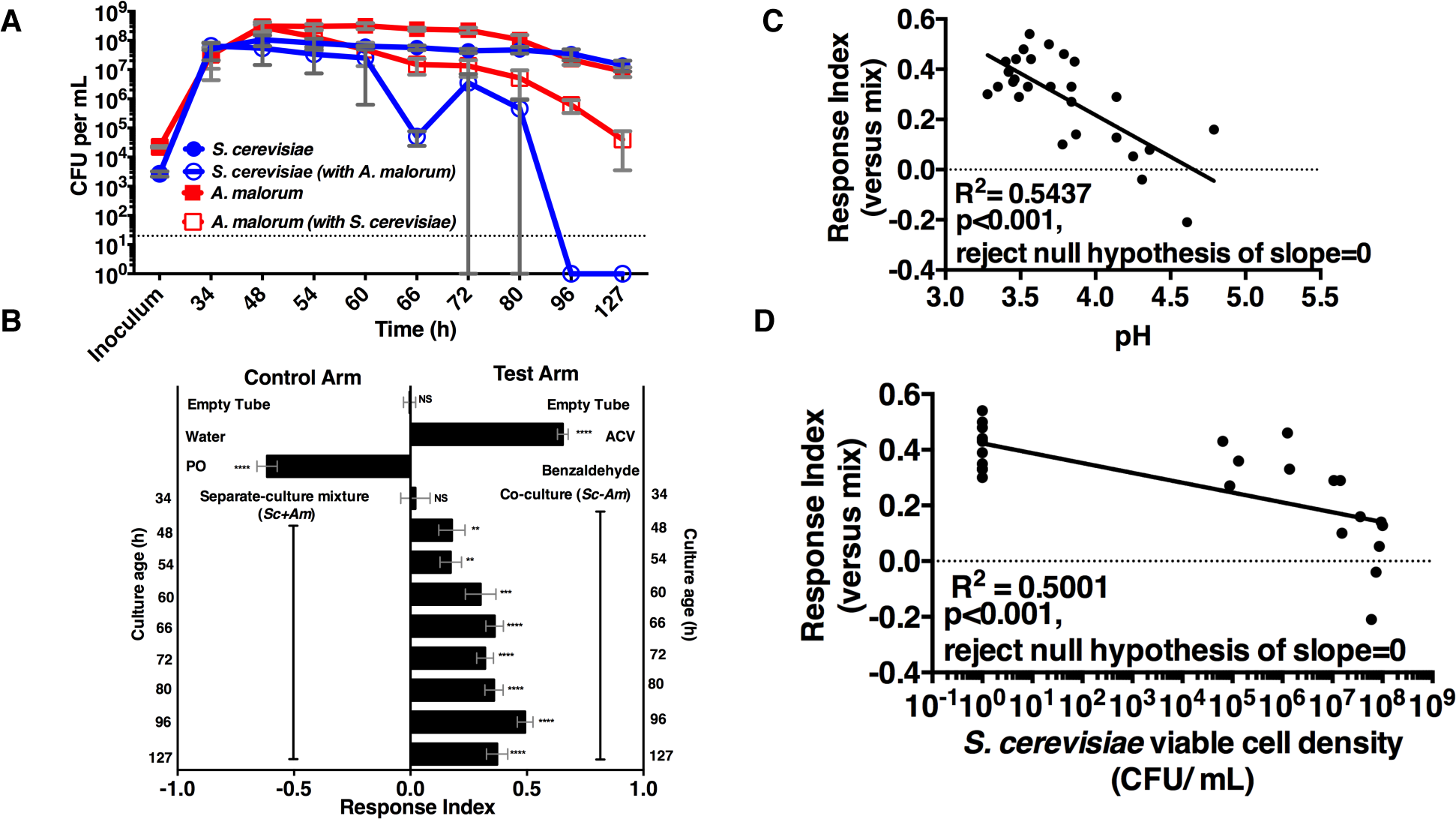
*Drosophila* temporal preference for metabolic exchange. **(A)** *S. cerevisiae* and *A. malorum* viable populations. Mean ± SEM of 2-3 experiments with one pooled replicate (2-3 cultures derived from the same colony) per experiment. Limit of detection is 20 CFU/mL. **(B)** *Drosophila* olfactory behavior toward co-cultured *S. cerevisiae* and *A. malorum* versus its separate-culture mixture as a function of culture age. Mean ± SEM of 16-18 replicates from three experiments. A t-test against a hypothetical value of zero Relationship between pH and *Drosophila* preference for the *S. cerevisiae* and *A. malorum* co-culture assessed deviance from 0. Symbols: NS p >0.05; * p ≤0.05; ** p ≤0.01; *** p ≤0.001; **** p ≤0.0001. **(C)** Relationship between pH and *Drosophila* preference for the *S. cerevisiae* and *A. malorum* co-culture versus the separate-culture mixture. Each data point represents the pH of a co-culture and the mean RI of *Drosophila* toward the same co-culture. A linear standard curve with an unconstrained slope was generated and compared to a null model with slope=0. The data fit to an unconstrained slope better than to the null model (p<0.0001, slope = -0.3295). **(D)** Relationship between *S. cerevisiae* populations and *Drosophila* preference for the co-culture versus the separate-culture mixture. Each data point represents viable *S. cerevisiae* populations of the culture along with the mean RI value toward the co-culture containing *S. cerevisiae*. A semilog standard curve with an unconstrained slope was generated and compared to a null model with slope=0. The data fit to an unconstrained slope better than to the null *Drosophila* preference for the co-culture versus the separate-culture mixture. Each data point represents model (p<0.0001, slope = -0.0349).

Mutants in broadly and narrowly tuned ionotropic and olfactory receptors (Irs and Ors, respectively, (45, 46)) were used to evaluate the role of *Drosophila* olfactory reception in discriminating the co-culture from the separate-culture mixture during and immediately following peak attraction (Figure 3-figure supplement 1). During the most attractive phase of the co-culture (e.g. 67-115 h), Or83b and Or42b showed significant attraction to the co-culture, whereas no role was detected for Irs or Or35a (Figure 3). As the co-culture proceeded (e.g. 139-163 h), attraction decreased and the role of Or42b and Or83b waned (Figure 3). Or83b is a required co-receptor for all other Or gene products (47) and Or42b, one of the most conserved olfactory receptors, detects esters and 1,1-diethoxyethane (48-50). These results suggest that Or42b enables *Drosophila* to distinguish the co-culture from the separate-culture mixture. Our results suggest that a non-Or83b factor explains ∼40% of *Drosophila* co-culture preference (Figure 3). Previous work found that Or83b is fully responsible for *Drosophila* attraction to apple cider vinegar (51), suggesting that the behavioral circuit activated by inter-species interactions of the co-culture is distinct from the circuit activated by apple cider vinegar.

**Fig. 3.**
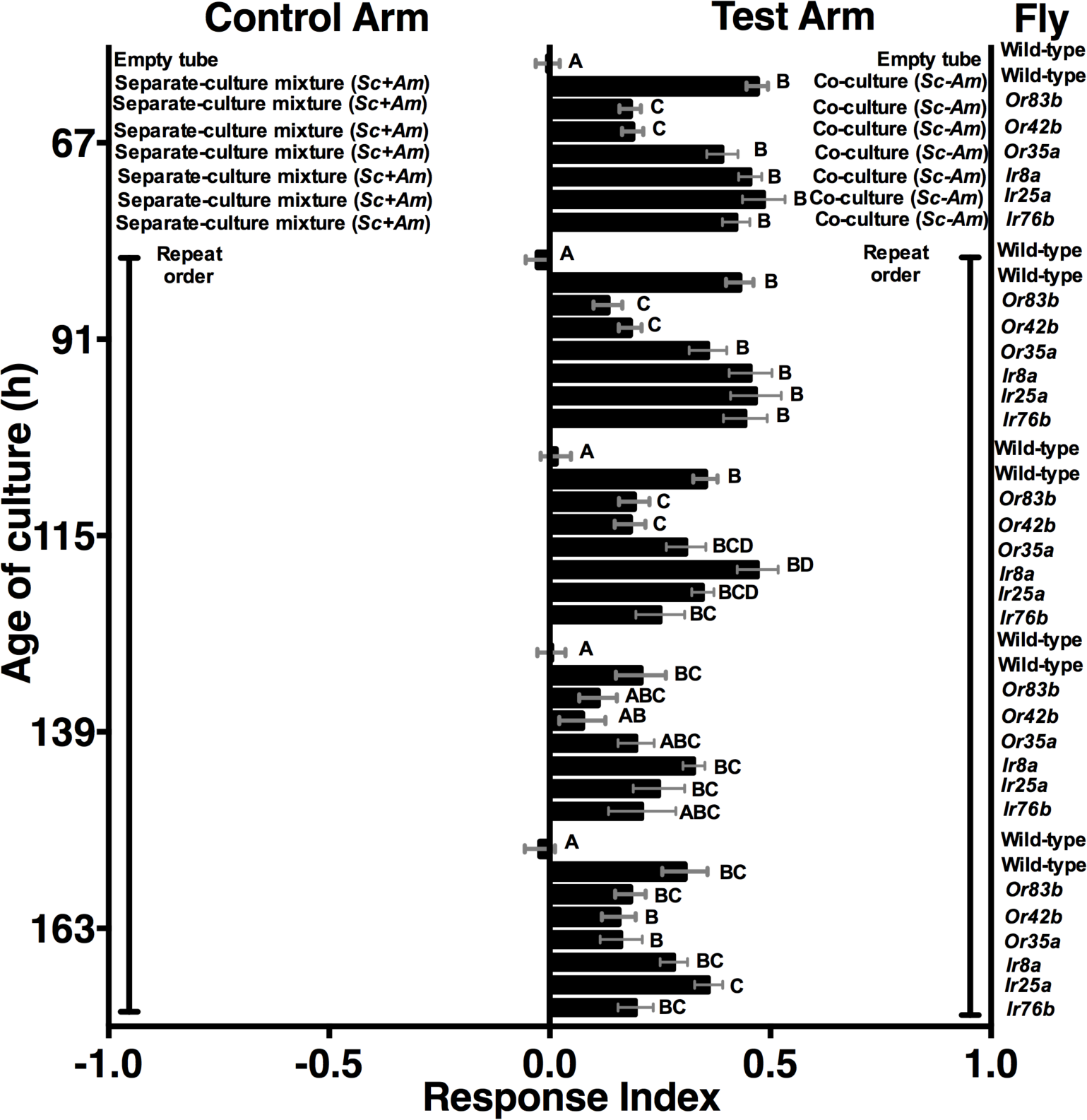
Role of olfactory receptor mutants in *Drosophila* detection of inter-species microbial interactions. Culture were grown at and after peak *Drosophila* attraction. Differences were assessed by a one-way ANOVA within each culture age. A Tukey’s multiple comparison test was used to assess significant differences between groups (p≤0.05), as indicated by unique letters. Mean +/- SEM of 12-24 replicates per time point per fly condition (n=2-4 experiments per time point).

We speculated that the emergent property of co-culture attractiveness might arise from a distinct metabolic profile of the co-culture. Using gas chromatography-mass spectrometry (GC-MS), we identified volatiles unique to or differentially produced in the co-culture compared to the separate-culture mixture. Five co-culture volatiles (ethanol, isobutanol, isoamyl alcohol, acetic acid, isoamyl acetate) were confirmed with standards (Table 1-figure supplements 1-2) and quantified with standard curves (Table 1-figure supplement 3). The alcohol concentrations were lower, and acetic acid and isoamyl acetate were unique in the co-culture relative to the other experimental groups (Table 1). The molecular profile was reminiscent of ethanol catabolism as the unique co-culture metabolic process. We therefore hypothesized that ethanol catabolism was the emergent metabolic process.

**Table 1.** Summary of volatiles detected using GC-MS. Relative abundance of volatiles in the co-culture (*S. cerevisiae* and *A. malorum grown* together) compared to the separate-culture mixture (*S. cerevisiae* and *A. malorum* grown separately, and their quantities added in during anaylsis). GC-MS captured volatiles with XAD-4 beads suspended above the cultures during growth; subsequently, beads were methanol-extracted (n=6 experiments, Table 1-figure supplements 1-2). Quantification is based on two experiments in which a linear regression was computed with standards (Table 1-figure supplement 3). Quantification is based on beads suspended above the cultures between 84 and 96 h of culture growth.

| *Identity* | *Standard confirmation* | *Relative quantification (co-culture: separate-culture mixture)* |
| --- | --- | --- |
| Ethanol | Y | 5.0-12.6-fold reduced |
| Isobutanol | Y | 7.3-24.7-fold reduced |
| Isoamyl acetate | Y | unique to co-culture |
| Isoamyl alcohol | Y | 3.6-6.4-fold reduced |
| Acetic acid | Y | unique to co-culture |

We next measured ethanol and acetic acid levels over time (24-156 h) and compared the chemical dynamics to *Drosophila* preference. Consistent with a relationship between ethanol catabolism, acetic acid anabolism, and *Drosophila* attraction, ethanol levels declined and acetic acid accumulated in the co-culture at a similar time of *Drosophila* co-culture preference (Figure 4A). Furthermore, as ethanol catabolism and acetic acid anabolism proceeded (36-96 h), *Drosophila* attraction toward the co-culture increased until 96 h, at which point it decreased, consistent with lower turnover of ethanol at the end of ethanol catabolism (Figure 4A, black line).

**Fig. 4.**
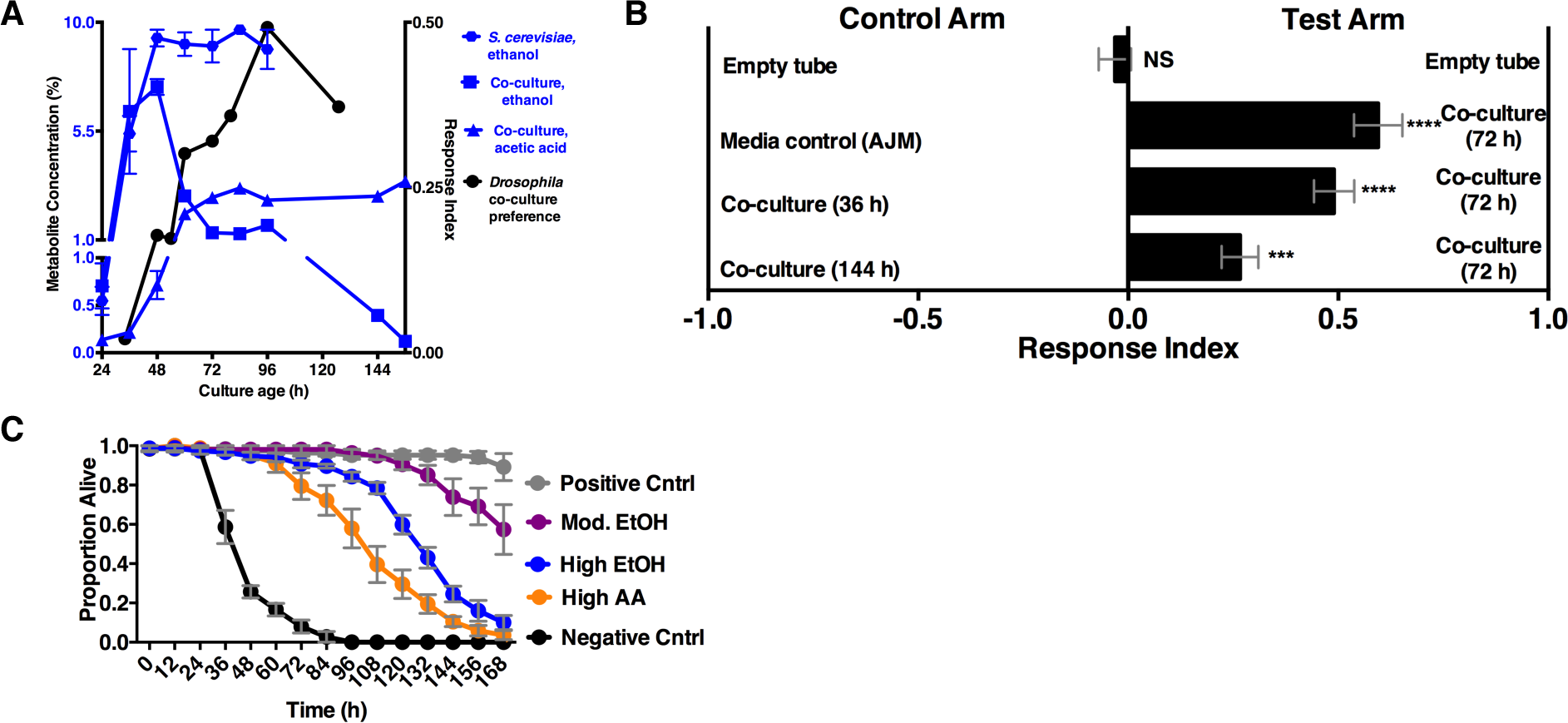
*Drosophila* behavior, survival, and ethanol catabolism. **A)** Dynamics of ethanol, acetic acid and *Drosophila* co-culture preference. Acetic acid was only detected in the co-culture. The abundance was of two values calculated from two experiments with three replicates per experiment (except acetic acid derived from a linear regression calculated from standards (figs. S6). Chemical data is the mean ± SEM of two values calculated from two experiments with three replicates per experiment (except acetic acid and ethanol concentrations at 144 and 156 h, which are from one experiment with three replicates). *Drosophila* co-culture preference is the mean value of the preference shown in Figure 2B. **B)** *Drosophila* preference for stages of ethanol catabolism. 72 h is ‘mid’ stage; 36 h is ‘early’ stage and 144 is ‘late’ stage. The co-culture contains *S. cerevisiae* and *A. malorum* grown for the time indicated. AJM= apple juice media. Data points represent the mean ± SEM of the combined results of two experiments with 8-10 total replicates per group. A t-test assessed whether the values were different from zero. C) *Drosophila* survival in the presence of acetic acid (AA), ethanol (EtOH) or the combination of the two in water. Groupings were based on concentrations of metabolites estimated from pre-ethanol catabolism (9.4% ethanol, High EtOH), middle-staged ethanol catabolism (1.4% ethanol, 2.8% acetic acid, Mod. EtOH) and post ethanol catabolism (3.42% acetic acid, High AA). Data represent mean +/- SEM of 5-6 replicates of one representative experiment of two. Mod. EtOH was significantly different from High EtOH condition from 108 h onward and High AA condition from 72 h onward (assessed by two-way ANOVA comparing time and condition, p≤0.05, Tukey’s correction). Negative Cntrl is water and Positive Cntrl is Shield’s & Sang M3 Insect Medium.

We hypothesized that *Drosophila* preferred the community during peak ethanol turnover (e.g. co-cultures at ∼72 h of growth) compared with the community pre-ethanol catabolism (e.g. co-culture at ∼36 h of growth) or during late-stage ethanol catabolism, in which ethanol turnover is low (e.g. co-culture at ∼144 h of growth; Figure 4A). Consistent with our hypothesis, *Drosophila* preferred the co-culture in the middle stage of ethanol catabolism to earlier or later stages of ethanol catabolism (Figure 4B).

We hypothesized that *Drosophila* preference for communities during peak ethanol turnover reflected fitness benefits derived from ingesting metabolites associated with different staged communities. To test this hypothesis, we measured *Drosophila* survival when given ethanol and acetic acid concentrations characteristic of microbial cultures at different stages of ethanol catabolism. *Drosophila* survival was highest when given metabolites associated with middle-staged ethanol catabolism compared with pre- or end-stage ethanol catabolism (Figure 4C). In sum, our results are consistent with *Drosophila* preference providing benefits associated with consumption of microbial community-generated metabolites.

To test directly whether ethanol catabolism underpinned *Drosophila* co-culture preference, we evaluated *Drosophila* preference for the co-culture harboring a mutant in *adhA*, which encodes pyrroloquinoline quinone-dependent alcohol dehydrogenase (PQQ-ADH-I), the enzyme that converts yeast-derived ethanol into acetaldehyde on path to acetic acid (34). Co-cultures using either *A. malorum* or *A. pomorum* wild-type (WT) along with *S. cerevisiae* were equally attractive to *Drosophila* (Figure 5-figure supplement 1). *Drosophila* preferred the co-culture containing *A. pomorum* WT versus a separate-culture mixture; however, *Drosophila* did not prefer the co-culture containing *A. pomorum adhA* versus a separate-culture mixture (Figure 5A), suggesting that ethanol catabolism is necessary for *Drosophila* to discriminate between the two conditions. *Drosophila* preferred the co-culture containing *A. pomorum* WT to the co-culture containing *A. pomorum adhA* (Fig. 5A). Moreover, *Drosophila* preferred to lay eggs in the co-culture containing *A. pomorum* WT to the co-culture containing *A. pomorum adhA*, further substantiating the importance of ethanol catabolism in *Drosophila* biology, as egg-laying decisions are crucial to reproductive success (Figure 5B).

**Fig. 5.**
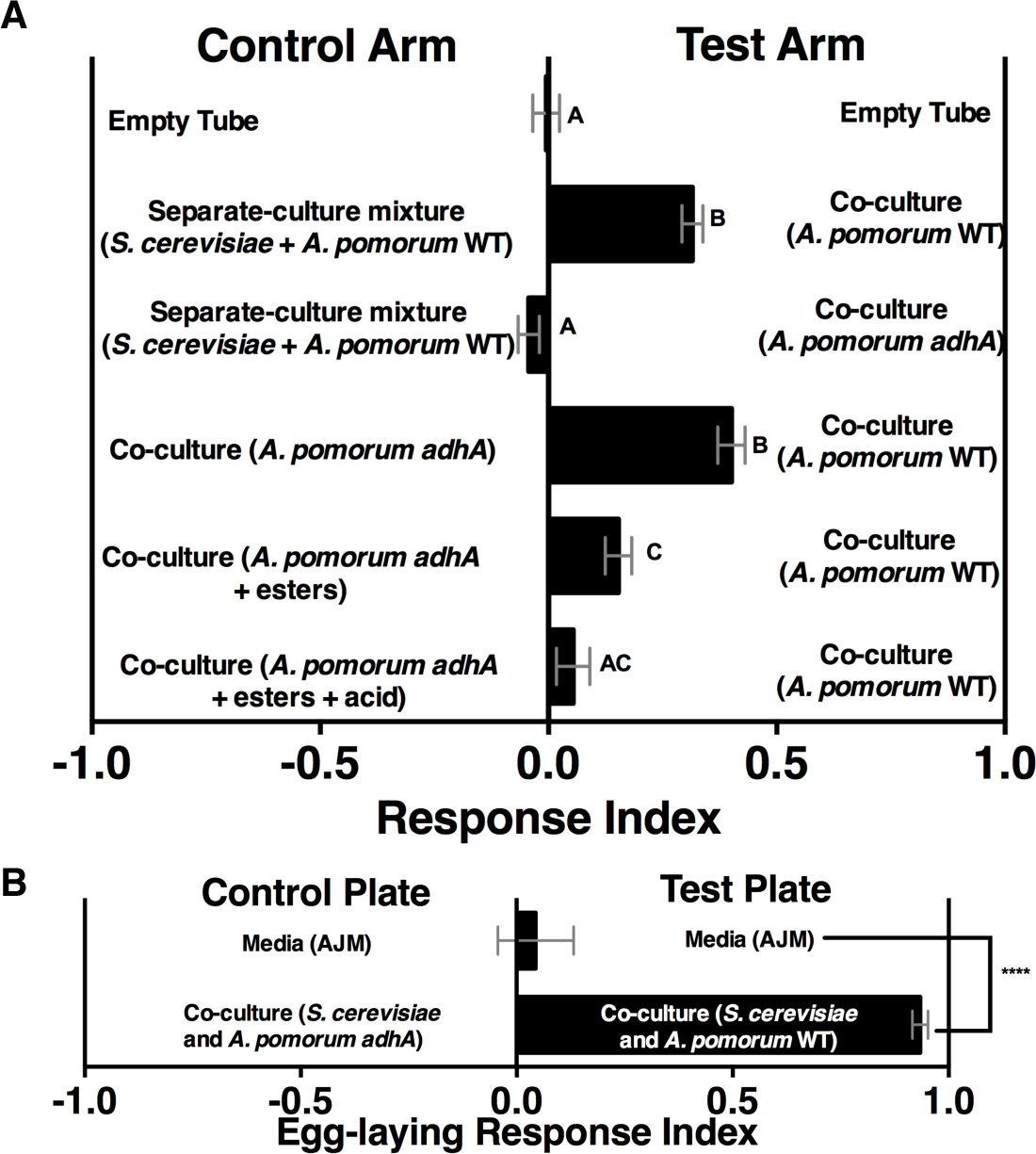
Role of ethanol catabolism in *Drosophila* olfactory and egg-laying behaviors. **(A)** *Drosophila* olfactory behavior toward specified conditions. Mean ± SEM of 2-7 experiments with 10-42 total replicates. Differences were assessed by a one-way ANOVA. A Tukey’s multiple comparison test was used to assess significant differences between groups (p≤0.05), as indicated by unique letters. Esters include ethyl acetate, isoamyl acetate, 2-phenethyl acetate, isobutyl acetate, 2-methylbutyl acetate, and methyl acetate; acid is acetic acid. Amounts added are based on physiological amounts in co-cultures and are found in Table 2. The co-culture contains *S. cerevisiae* and the specified *A. pomorum* strain. acid= acetic acid **(B)** *Drosophila* was given a choice to lay eggs in a co-culture containing *S. cerevisiae* and *A. pomorum* wild-type (WT) or *S. cerevisiae* and *A. pomorum adhA*. The co-cultures were grown for 96 h and mixed 1:1 with a 1.6% agarose solution. *Drosophila* was allowed to lay eggs for eight hours. Difference between the two groups was assessed by an unpaired t-test (**** p≤0.0001). Mean ± SEM of 16-18 replicates from two experiments.

We next identified additional metabolites unique to the co-culture using solid-phase microextraction gas chromatography-mass spectrometry (SPME GC-MS). Acetic acid, six acetate esters, an acetaldehyde metabolic derivative (acetoin), a putative acetaldehyde metabolic derivative (2,4,5-trimethyl-1,3-dioxolane), and two unknown metabolites were more abundant in the co-culture relative to the separate-culture mixture or co-culture with *A. pomorum* lacking *adhA* (Table 2, Table 2-figure supplements 1-2). To determine the molecular basis for *Drosophila* co-culture preference, select metabolites were added to the co-culture containing *A. pomorum adhA*. Esters and acetic acid, but not esters alone, were sufficient to fully restore *A. pomorum adhA* attractiveness to *A. pomorum* WT levels (Figure 5A).

**Table 2.**
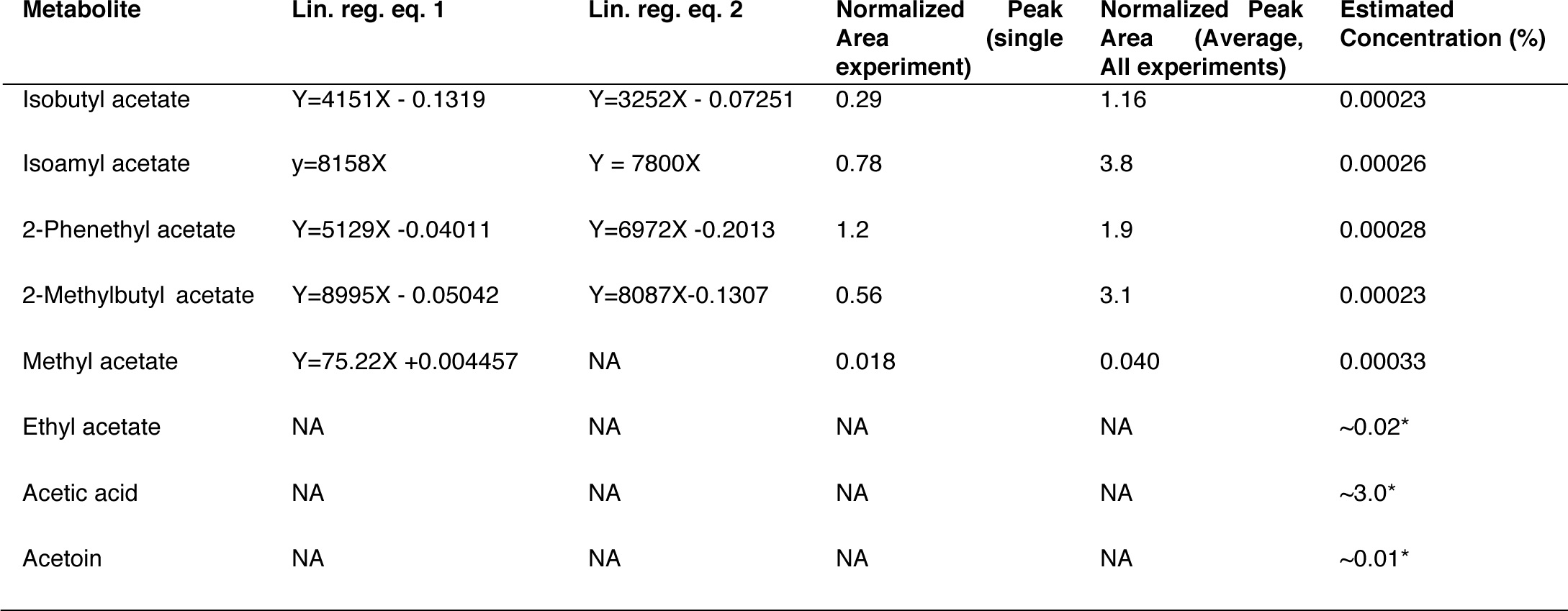
Estimated concentrations of key metabolites in the co-culture using SPME GC-MS. Estimated concentrations of differentially concentrated or unique metabolites in the co-culture. Linear regression equations (Lin. reg. eqs. 1 and 2) were estimated from individual experiments in which peak areas of different concentrations of metabolites were fitted with a linear regression (Table 2-figure supplement 2). Normalized peak areas correspond to the specified metabolites in co-cultures containing *S. cerevisiae* and *A. malorum*. Separate estimates were derived from a normalized peak area estimated from a single experiment (co-culture and standard samples were from a run with similar internal standard signal) or from the mean normalized peak area estimated from all experiments (co-cultures were run over four days, standards were run on two days). The final estimated concentration was an average of all estimated concentrations (n=4 estimates (two from each standard regression equation times two estimates of the normalized peak area), execpt for methyl acetate, n=2 estimates). The estimated concentrations (exept acetoin) were added to the co-culture containing *A. pomorum adhA* (Figure 5A). *Ethyl acetate, acetic acid and acetoin concentrations were estimated from standards in Table 2-figure supplement 1.

Although acetate and its metabolic derivatives were sufficient for *Drosophila* co-culture preference, acetaldehyde is a reactive intermediate during ethanol catabolism whose metabolic derivatives might be increased in microbial communities compared with individual microbial cultures. Consistent with this idea, acetoin was moderately increased in the co-culture compared with the separate-culture mixture (Table 2-figure supplement 1); strikingly, acetoin was increased ∼27-fold in the tri-culture (*S. cerevisiae*, *A. malorum*, and *L. plantarum*) compared to the co-culture (Figure 6A, B, Figure 6-figure supplement 1) and was attractive to *Drosophila* (Figure 6-figure supplement 2). In sum, emergent metabolites from 2- and 3-membered communities, including acetaldehyde metabolic derivatives, attract *Drosophila*.

**Fig. 6.**
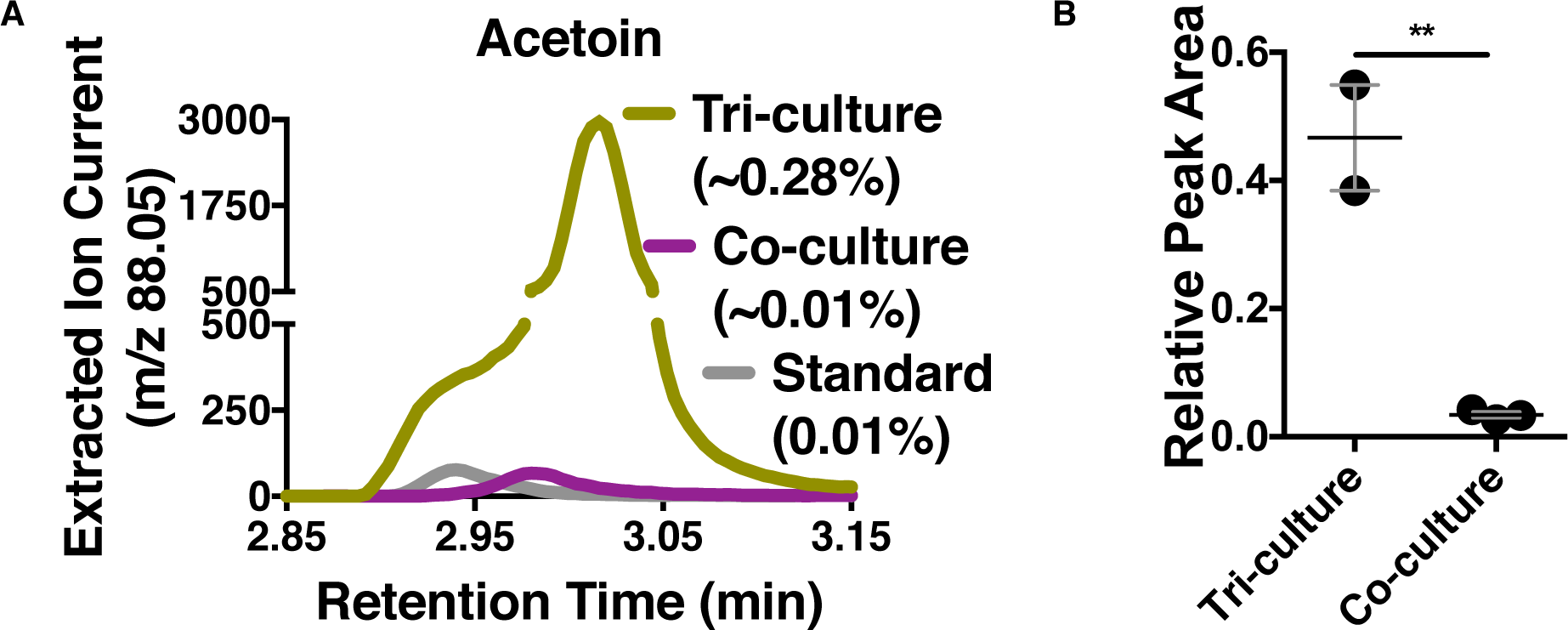
Acetoin as an attractive metabolite increased in the 3-membered communitiys **A)** Representative chromatogram of the tri-culture (*S. cerevisiae-A. malorum*-*L. plantarum*) compared to the co-culture (*S. cerevisiae* and *A. malorum)*. **B)** Estimated quantification of is based on a linear regression of acetoin (figure 6-figure supplement 1). Relative quantification of acetoin in the tri-culture (one replicate with *A. malorum* and one replicate with *A. pomorum* from separate days) and the co-culture (one replicate with *A. malorum* and two replicates with *A. pomorum* from separate days). Difference in peak areas was assessed by an unpaired two-tailed t-test (** p≤0.01).

To investigate further the potential role of acetaldehyde and its metabolic derivatives in *Drosophila* behavior, we performed a dose response in which acetaldehyde was added to the separate-culture mixture (Figure 6-figure supplement 3A) to evaluate its ability to induce attractiveness to co-culture levels. Even at the lowest tested levels, acetaldehyde supplementation stimulated the separate-culture mixture to attractiveness levels equal to the co-culture (Figure 6-figure supplement 3A); moreover, acetaldehyde and its metabolic derivatives ‘over-restored’ preference of the separate-culture mixture beyond WT co-culture levels at its most attractive dose (Figure 6-figure supplement 3A). Finally, three acetaldehyde metabolic derivatives—acetoin, 1,1-diethoxyethane (an acetal), and 2,3-butanedione—were sufficient to induce the attractiveness of the separate-culture mixture to WT co-culture levels using concentrations of each metabolite at or below the physiological concentration of acetoin found in the tri-culture (Figure 6-figure supplement 3B). Creating a pure metabolite mixture comprised of key metabolic groups produced by microbial communities and identified in this study (esters, acetaldehyde metabolic derivatives, alcohols, acid) attracted *Drosophila* similarly to the co-culture (Figure 6-figure supplement 4A, B). Interestingly, the acetaldehyde metabolic derivatives alone were sufficient and the most attractive (Figure 6-figure supplement 4C, D), supporting acetaldehyde metabolic derivatives as potent *Drosophila* attractants.

Overall, our results suggest that both esters and acetaldehyde metabolic derivatives are keystone microbial community metabolites that attract *Drosophila*. We next created a simple 9-metabolite mixture in water (containing only 1 acid, 4 esters, and 4 acetaldehyde metabolic derivatives) and measured *Drosophila* preference toward this mixture in relation to the yeast-acetic acid bacteria co-culture, the yeast-acetic acid bacteria-lactic acid bacteria microbial community, or apple cider vinegar (ACV). The defined mixture used concentrations for each acetaldehyde metabolic derivative similar to the concentration of acetoin in the tri-culture and ester and acid concentrations that were in the range of detected in the co-culture. The defined 9-metabolite mixture was more attractive than all other conditions (Figure 6-figure supplement 5). In sum, acetaldehyde metabolic derivatives and esters are potent *Drosophila* attractants whose detection may signal the presence of actively metabolizing, multispecies microbial communities.

## Discussion

Here, we have demonstrated how emergent properties of a microbial community—volatile profile, population dynamics, and pH—influence *Drosophila* attraction, survival, and egg-laying behaviors. Our study is the first to identify the consequences of microbe-microbe metabolic exchange on animal behavior and discovers additional microbial interactions for further mechanistic study (Figure 1D).

Microbe-microbe metabolic exchange generates unique and quantitatively different volatiles from those resulting from individual microbial metabolism (Tables 1 & 2, Figure 7). *Acetobacter*-generated acetate coupled to *Saccharomyces*-derived alcohols spawn diverse acetate esters (Table 1 & Table 2). We hypothesize that more complex and diverse communities, comprising alcohol-producing yeasts, acetate-producing *Acetobacter*, and lactate-producing *Lactobacillus*, will generate a wider array of attractive esters (Figure 7). The community of *S. cerevisiae*, *A. malorum*, and *L. plantarum* emitted higher levels of acetoin and attracted *Drosophila* more strongly than the co-culture of *S. cerevisiae* and *A. malorum* (Figure 1D, Figure 6). Acetoin and 2,3-butanedione are formed by an α-acetolactate intermediate in bacteria and directly from acetaldehyde in yeast (52). We therefore hypothesize that communities of yeasts and bacteria may emit high levels of attractive acetaldehyde metabolic derivatives (Figure 7).

**Figure 7.**
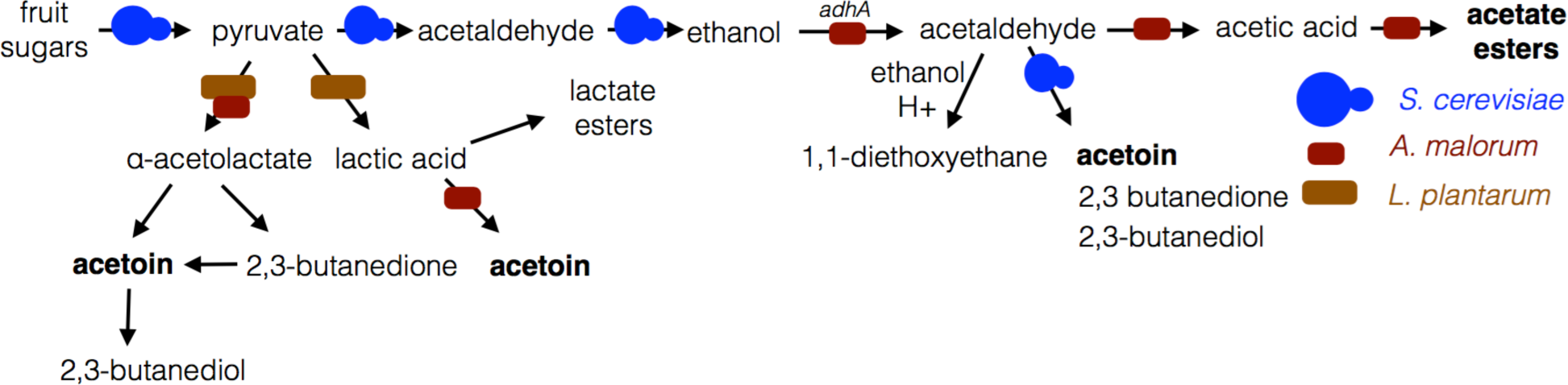
Model of microbe-microbe metabolite exchange. Bolded are metabolites increased due to microbe-microbe interactions.

*Drosophila* behavioral studies have mostly focused on yeasts, even though *Drosophila* evolved in a bacterial-rich environment. Our results suggest that non-yeast microorganisms, especially when grown in microbial communities, affect *Drosophila* behaviors. We reason that additional investigations that couple chemical microbial ecology with *Drosophila* behavior will herald the discovery of microbe-influenced behaviors and microbial community-generated metabolites.

This study demonstrates the coordination of ethanol synthesis and catabolism by *S. cerevisiae* and *Acetobacter,* respectively, and the role of ethanol in *Drosophila* behavior and survival. Non-*Saccharomyces Drosophila* microbiome members also make ethanol (53) and diverse acetic acid bacteria catabolize ethanol, generalizing our findings to other microbial community combinations. Ethanol can have deleterious or beneficial fitness consequences for *Drosophila* depending on concentration (54, 55) and ecological context (56). Our results show that *Drosophila* uses products of inter-species microbiome metabolism to detect a community that titrates ethanol concentration optimally for the host. Work that further dissects the consequences of acetic acid and ethanol concentrations on *Drosophila* biology and investigates other community-level metabolic profiles will be of interest to enrich the chemical and ecological portrait of the *Drosophila* microbiome.

Our work raises questions about the consequences of the observed behavior on microbiome assembly and stability in the *Drosophila* intestine. *Drosophila* possesses specific and regionalized gut immune responses to the microbiome (57-60) implying a tolerant environment in which privileged microbiome members are maintained and reproduce in the *Drosophila* intestine. Other work suggests that *Drosophila* acquires its adult microbiome from exogenous sources, that adult microbiome abundance drops without continuous ingestion of exogenous microorganisms, and that the microbiome can be shaped by diet (19, 25, 26). As such, a combination of internal mechanisms, exogenous factors, and host behavior likely sculpt the microbiome; determining the relative contribution of each will be important moving forward. Complicating our understanding of the contribution of these factors is the opaque distinction between ‘microbiome’ and ‘food’, since both are ingested from the environment (61). To dissect the formation and stability of the *Drosophila* microbiome, the fate of ingested microorganisms needs to be monitored and microbial intestinal reproduction needs to be surveyed as a function of *Drosophila* behavior, age, immune status, microbiome membership, and nutritional state (e.g. using synthetic diets without yeast; (34, 62)).

In sum, our results support a model in which the *Drosophila* olfactory system is tuned to fruity (e.g., esters) and buttery (several acetaldehyde metabolic derivatives, such as 2,3-butanedione) smelling metabolites promoted by microbe-microbe interactions. We anticipate that accounting for microbial interactions in diverse host-microbe studies will lead to new insights into diverse aspects of microbial-animal symbioses.

## Acknowledgements

We thank Dr. Fabian Staubach (Stanford University, USA), Dr. Angus Chandler (University of California, Berkeley, USA), Dr. Matthew Goddard (The University of Auckland, New Zealand), Dr. Dan Jarosz (Stanford University, USA) and the Phaff Yeast Culture Collection (UC Davis, USA) for microbial strains. We thank Dr. Ryan Joseph and Dr. Karen Menuz (Yale University/University of Connecticut) for help with the *Drosophila* behavioral experiments; the Yale West Campus Analytical Core for use of the GC-MS; Dr. Terence Wu and Dr. Eric Patridge (Yale University, USA) for advice and experimental help with the GC-MS; Dr. Scott Strobel and Michelle Legaspi for use of and help with SPME GC-MS; Dr. John Carlson (Yale University, USA) for the T-maze and several *Drosophila* lines. We also thank Dr. Won Jae Lee (Seoul National University, S. Korea) for the *A. pomorum* WT and *adhA* strains, and John Carlson, Craig Crews, and Andrew Goodman (Yale University, USA) for helpful discussions. This work was supported by National Institutes of Health Grants NIDDKRC12T32GM007499-36, 5T32HG003198-10, 1R01GM099563, 7RC1DK086831. The raw data for the metabolomics experiments can be found at https://figshare.com/account/home#/projects/4735.

## SUPPLEMENTARY MATERIALS

**Table S1.**
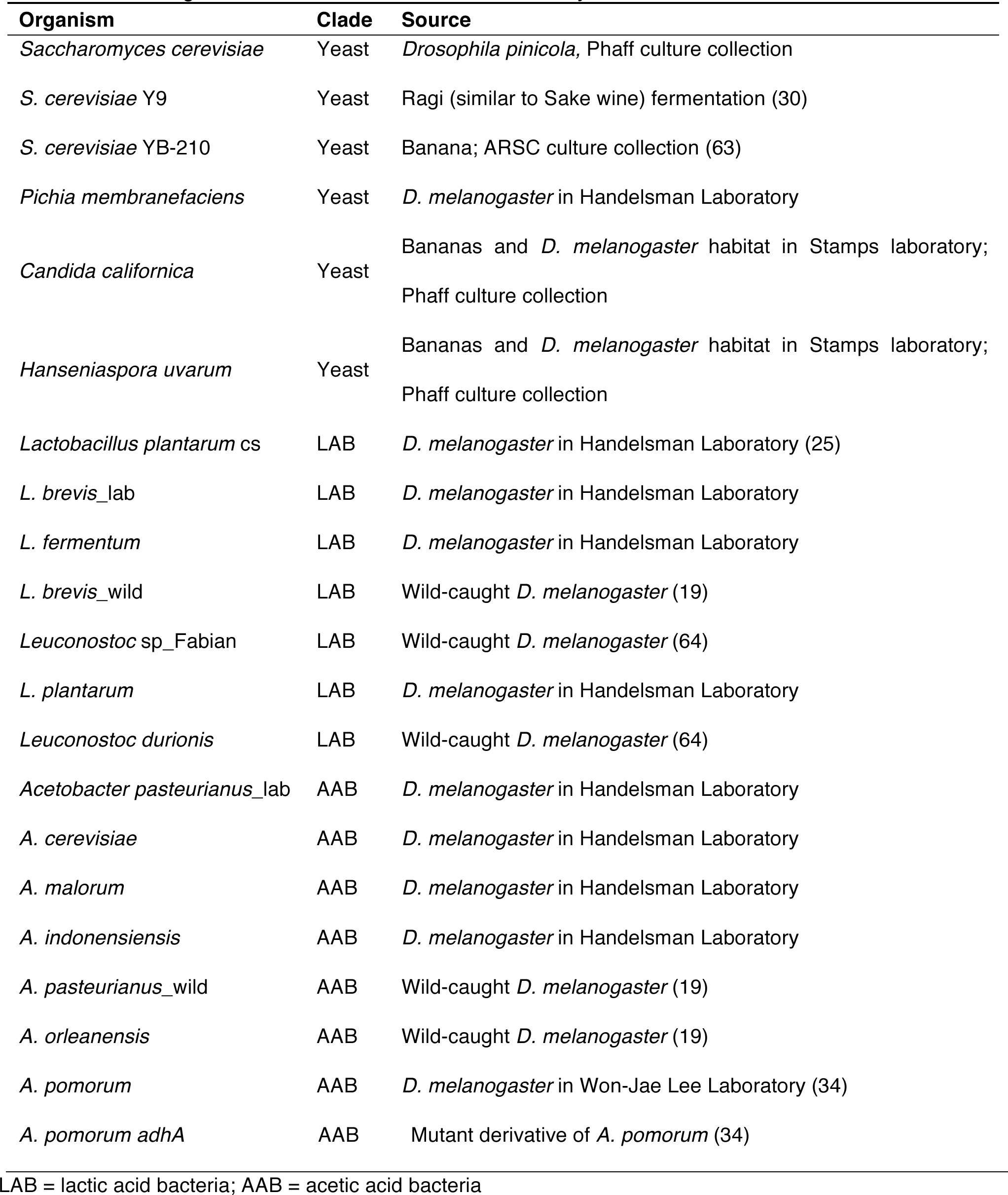
Microorganisms and their sources used in this study.

**Figure 1-figure supplement 1.**
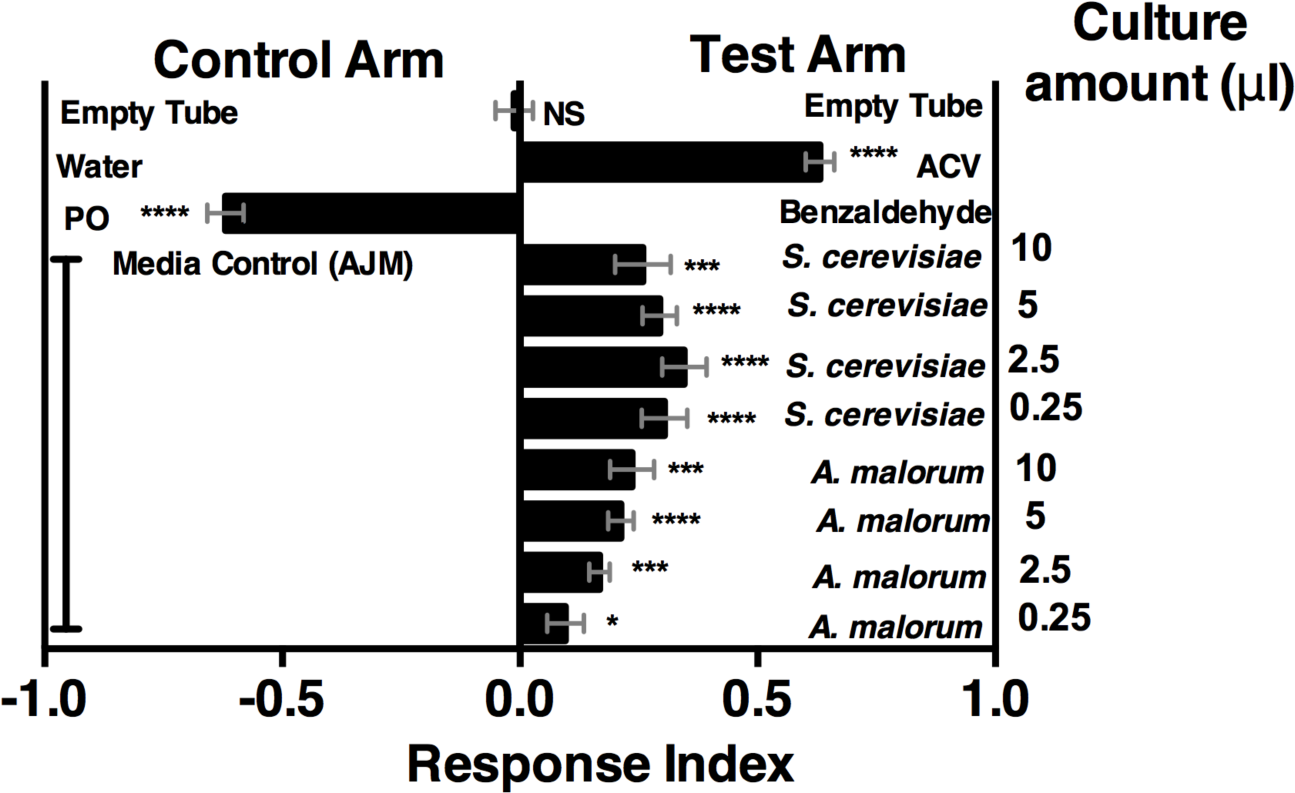
*Drosophila melanogaster* olfactory behavior toward different culture volumes of *Saccharomyces cerevisiae* and *Acetobacter malorum*. The top three experimental groups are controls: Mock (empty tube versus empty tube) recapitulates alternating of test and control arms, as in all experimental groups; apple cider vinegar (ACV (25% in water)) is the positive control and tested against water only; Benzaldehyde (1%) is the negative control and is a 100-fold dilution of benzaldehyde in paraffin oil (PO) tested against paraffin oil only. In all experimental groups, 10µl of total volume was used; the culture amount is specified, when appropriate, on the right-hand portion of the plot. The remaining volume in the microbial groups is water. Media control (AJM) is always 5µl of AJM mixed with 5µl of water. Data points represent the mean +/- SEM combined from three experiments (n=12 per experimental group). A t-test was performed against a hypothetical value of 0 to test for significance and is denoted by the following system (consistent throughout manuscript): NS p >0.05; * p ≤0.05; ** p ≤0.01; *** p ≤0.001; **** p ≤0.0001. Based on this analysis, we used a total microbial culture volume of 5 µl throughout the manuscript, unless otherwise noted.

**Figure 2-figure supplement 1.**
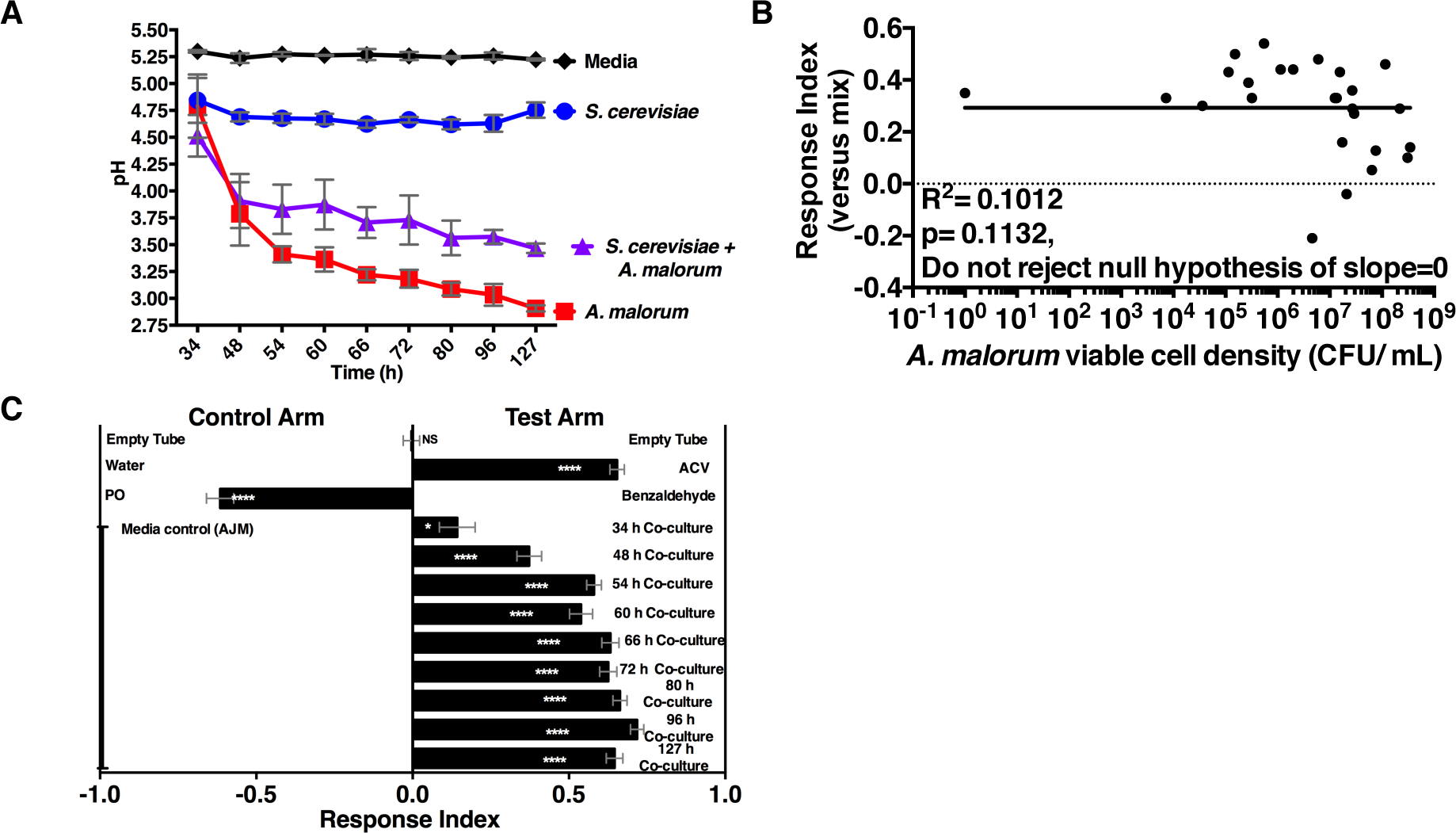
Properties of the co-culture and its relationship to *Drosophila* preference **(A)** pH of experimental groups as a function of microbial growth time. Mean pH ± SEM of three experiments with one pooled replicate per experiment. **(B)** Relationship between *A. malorum* populations and *Drosophila* preference for the co-culture versus the separate-culture mixture. Each data point represents viable *A. malorum* populations in the co-culture along with the mean RI value toward the co-culture containing *A. malorum*. A semi-log standard curve with an unconstrained slope was generated and compared to a null model with slope=0. The data do not fit to an unconstrained slope better than to the null slope= 0 model (p=0.1132). **(C)** *Drosophila* attraction to the co-culture versus sterile media as a function of co-culture age (grown 34 h – 127 h). Mean RI ± SEM of three experiments with 16-18 total replicates. T-test against a hypothetical value of 0 was performed. NS p >0.05; * p ≤0.05; ** p ≤0.01; *** p ≤0.001; **** p ≤0.0001 Significance denoted beside or within bars of each experimental group. ACV = apple cider vinegar (25% in water); PO = paraffin oil.

**Figure 3-figure supplement 1.**
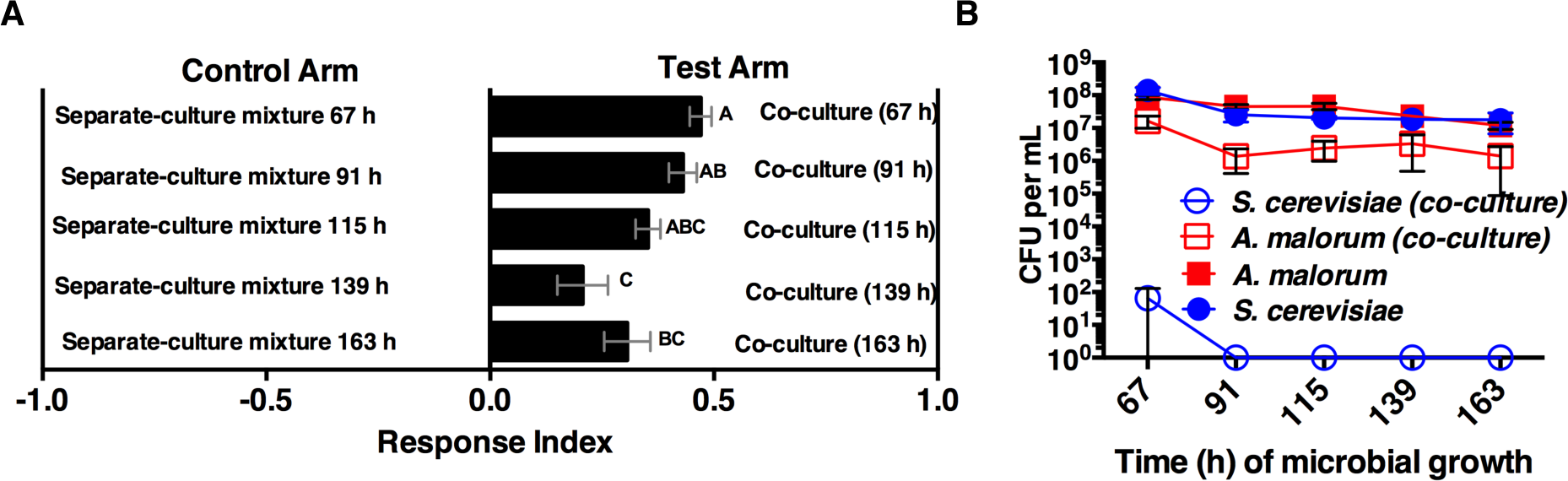
**A)** Attraction of wild-type *Drosophila* to different aged co-cultures (grown 67-163 h, *S. cerevisiae* and *A. malorum*). Mean ± SEM of 12-24 replicates per group (n=2-4 experiments). One-way ANOVA assessed significance between groups. A Tukey’s multiple comparison test was used to assess significant differences between groups (p≤0.05), as indicated by unique letters. **B)** Corresponding viable counts at different times of microbial growth. Mean ± SEM of 2-3 replicates.

**Table 1-figure supplement 1.**
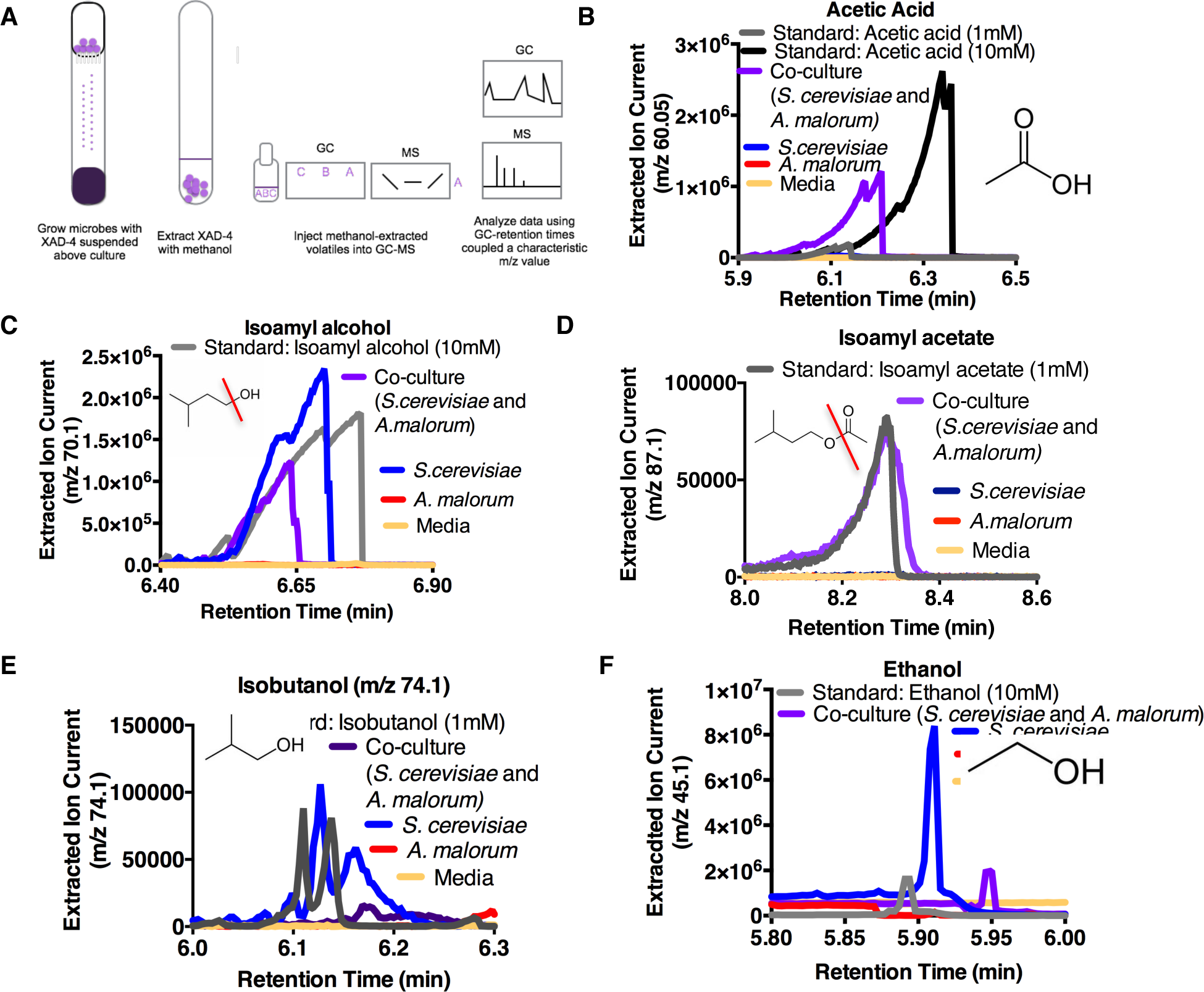
Extracted ion chromatograms of the five metabolites detected by gas chromatography-mass spectrometry (GC-MS). **(A)** Schematic depicting the experimental setup **(B-F)** Representative extracted ion chromatograms from one replicate (out of three total) of one experiment (out of 3-4 total) of m/z values corresponding to major metabolites spectrometry (GC-MS). **(A)** Schematic depicting the experimental setup **(B-F)** Representative extracted ion chromatograms acetate (**D)** isobutanol **(E)**, and ethanol **(F)** were identified as the five major metabolites in the co-culture (*S. cerevisiae* and identified in the experimental conditions along with appropriate standards. Acetic acid **(B)**, isoamyl alcohol **(C)**, isoamyl grown alone. Extracted ion chromatograms were constructed using the m/z value in the title of each graph. For acetic acid *A. malorum*). Isoamyl alcohol **(C)**, ethanol **(E),** and isobutanol **(F)** were identified as the major metabolites in *S. cerevisiae* corresponds to the molecular weight minus one (hydrogen). For isoamyl alcochol, the m/z used corresponds to the loss of and isobutanol, the m/z value used corresponds to the molecular weight of the molecule. For ethanol, the m/z used value corresponds to the molecule shown within the graph. In all cases, figures showing the complete mass spectra the hydroxyl group (depicted), which may have picked up a hydrogen and been lost as water. For isoamyl acetate, the m/z value corresponds to the molecule shown within the graph. In all cases, figures showing the complete mass spectra between the metabolite and standard are found in Table 1-figure supplement 2. Microorganisms were grown 72-96 hours.

**Table 1-figure supplement 2.**
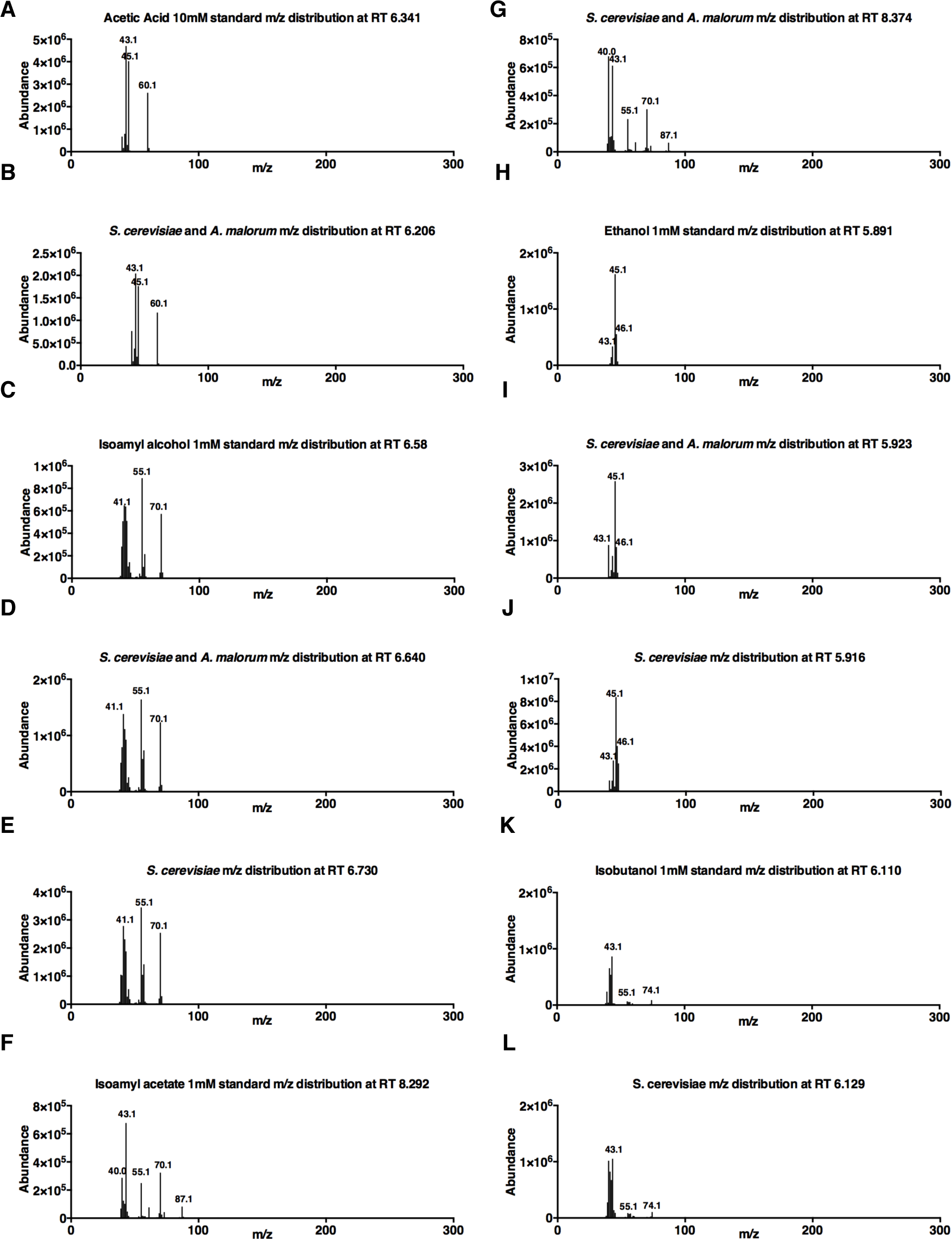
Representative spectra of acetic acid (**A-B)**, isoamyl alcohol (**C-E)**, isoamyl acetate (**F-G)**, ethanol **(H-J)** and isobutanol **(K-L)** in standard and experimental samples. Standard concentrations are denoted on individual graphs. All mass spectra are one replicate (out of 3-4 experiments with three replicates per experiment).

**Table 1-figure supplement 3.**
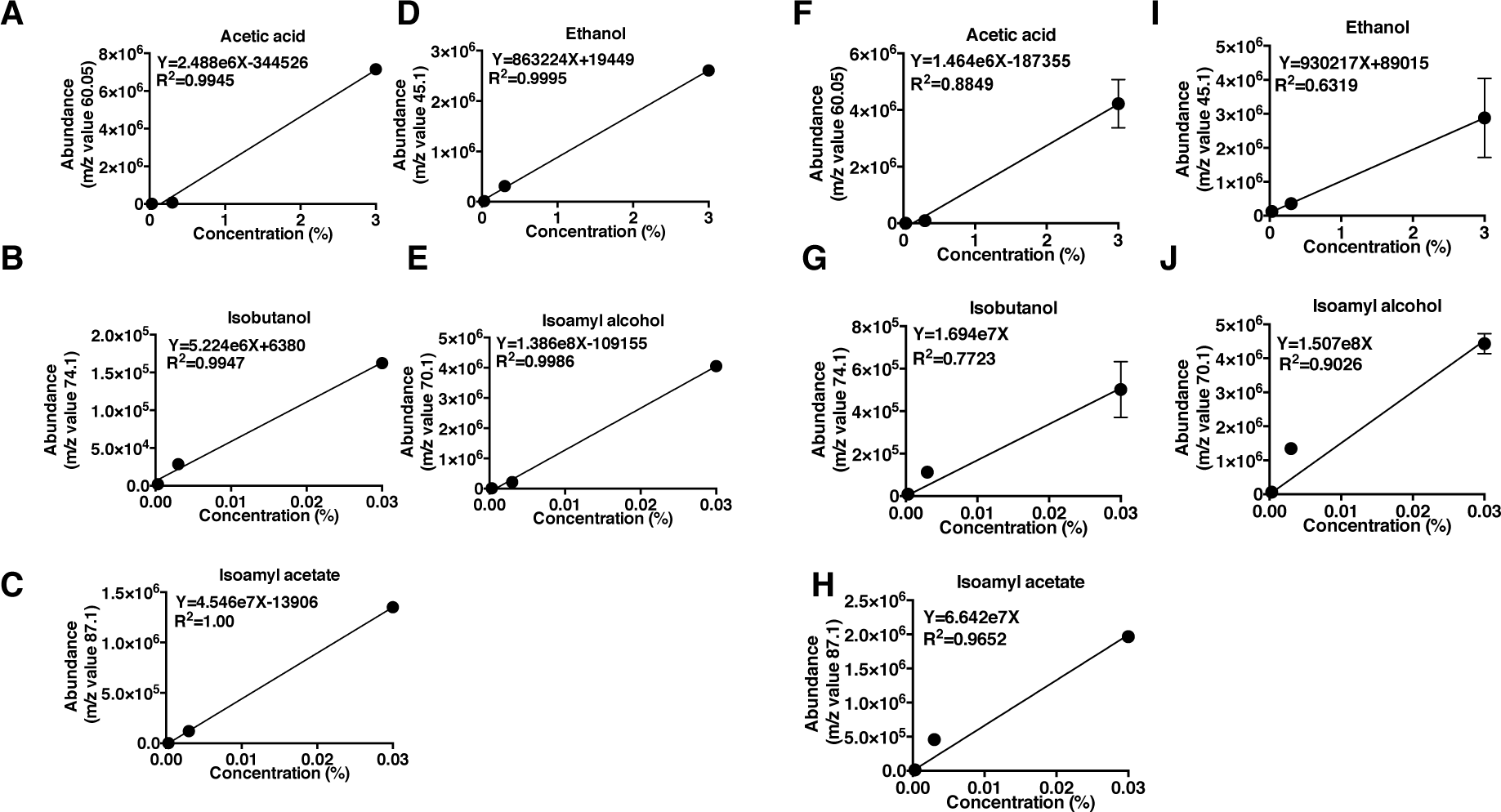
Estimation of volatile quantity using GC-MS. Separate experiments are graphed in panels **(A-E)** and **(F-J)**. **(A-E)** Data points represent the value of a single replicate per concentration for each standard. The abundance of a single m/z value at a specific retention time was chosen for each standard. The values were fitted with a linear regression and the equation was used to estimate the concentration of the five metabolites in the experimental samples from the same experiment. (**F-J)** Data points represent the mean ± SEM of three replicates for a given concentration for each standard. The abundance of a single m/z value at a specific retention time was chosen for each standard. The values were fitted with a linear regression. The equation was used to estimate the concentration of the 5 metabolites in the experimental samples from the same experiment. When applicable an equation was calculated when the line was forced to go through X,Y=0,0; these equations were used to calculate the concentrations of isoamyl alcohol, isoamyl acetate, and isobutanol.

**Figure 5-figure supplement 1.**
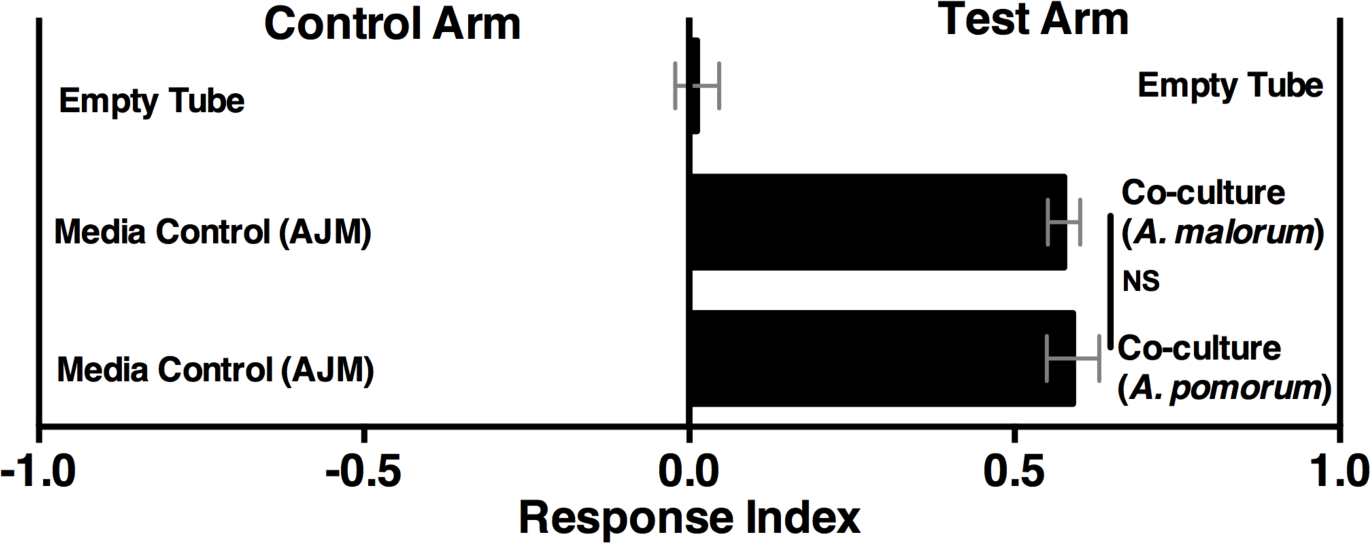
*Drosophila* behavior toward the co-culture using *A. malorum* or *A. pomorum. Drosophila* behavior toward co-cultures grown for 96 h using *A. malorum* or *A. pomorum* versus a media control (AJM = apple juice medium). Result of two experiments with six replicates each. Data points represent mean ± SEM. An unpaired two-tailed t-test assessed whether *Drosophila* behavior toward the co-cultures was different (NS p > 0.05).

**Table 2-figure supplement 1.**
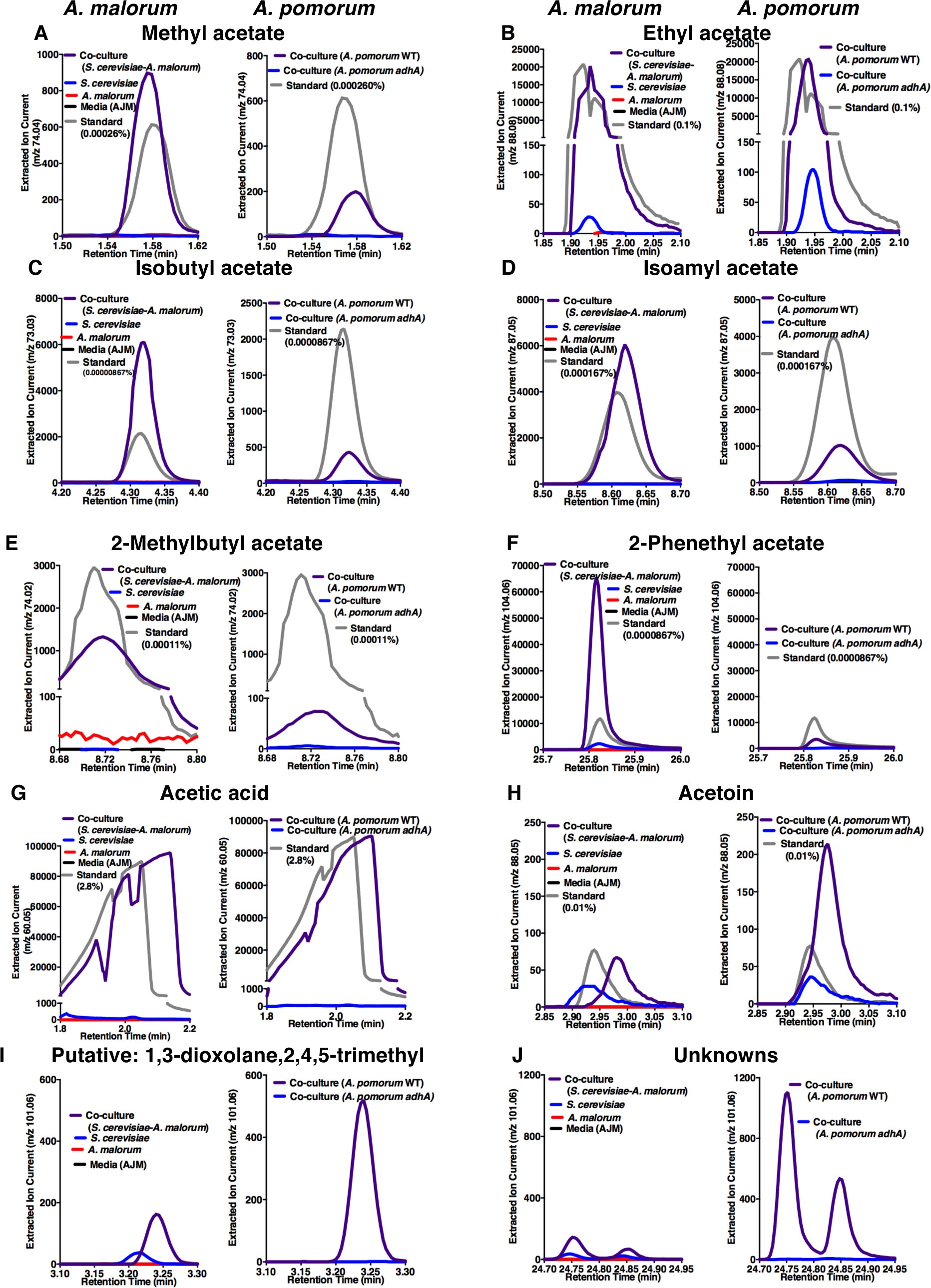
Extracted ion chromatograms of differentially emitted or unique metabolites in the co-culture according to solid phase microextraction gas chromatography-mass spectrometry (SPME GC-MS). Specific metabolites are displayed above each panel. For each panel, the left-most plot compares the co-culture containing *S. cerevisiae* and *A. malorum* to *S. cerevisiae* grown alone, *A. malorum* grown alone, or media (AJM (apple juice medium)); the right-most plot compares the co-culture containing *S. cerevisiae* and *A. pomorum* wild-type to the co-culture containing *S. cerevisiae* and *A. pomorum adhA*, since *A. pomorum adhA* is required for *Drosophila* co-culture preference (Figure 5A). The two plots within the same panel contain the same standard. The y-axis for each plot is the ion current for a m/z value that discriminates the metabolite of interest over a specific retention time window. The following m/z values were chosen for each metabolite based on standards or, in the cases of putative and unknown metabolites (**I** and **J)** were chosen from the experimental groups: **(A)** m/z 74.04 **(B)** m/z 88.08 **(C)** m/z 73.03 **(D)** 87.05 **(E)** 74.02 **(F)** 104.04 **(G)** 60.05 **(H)** 88.05 **(I)** 101.06 **(J)** 101.06. Each panel is one representative replicate of 1 experiment (out of 3-5 total replicates in three experiments).

**Table 2-figure supplement 2.**
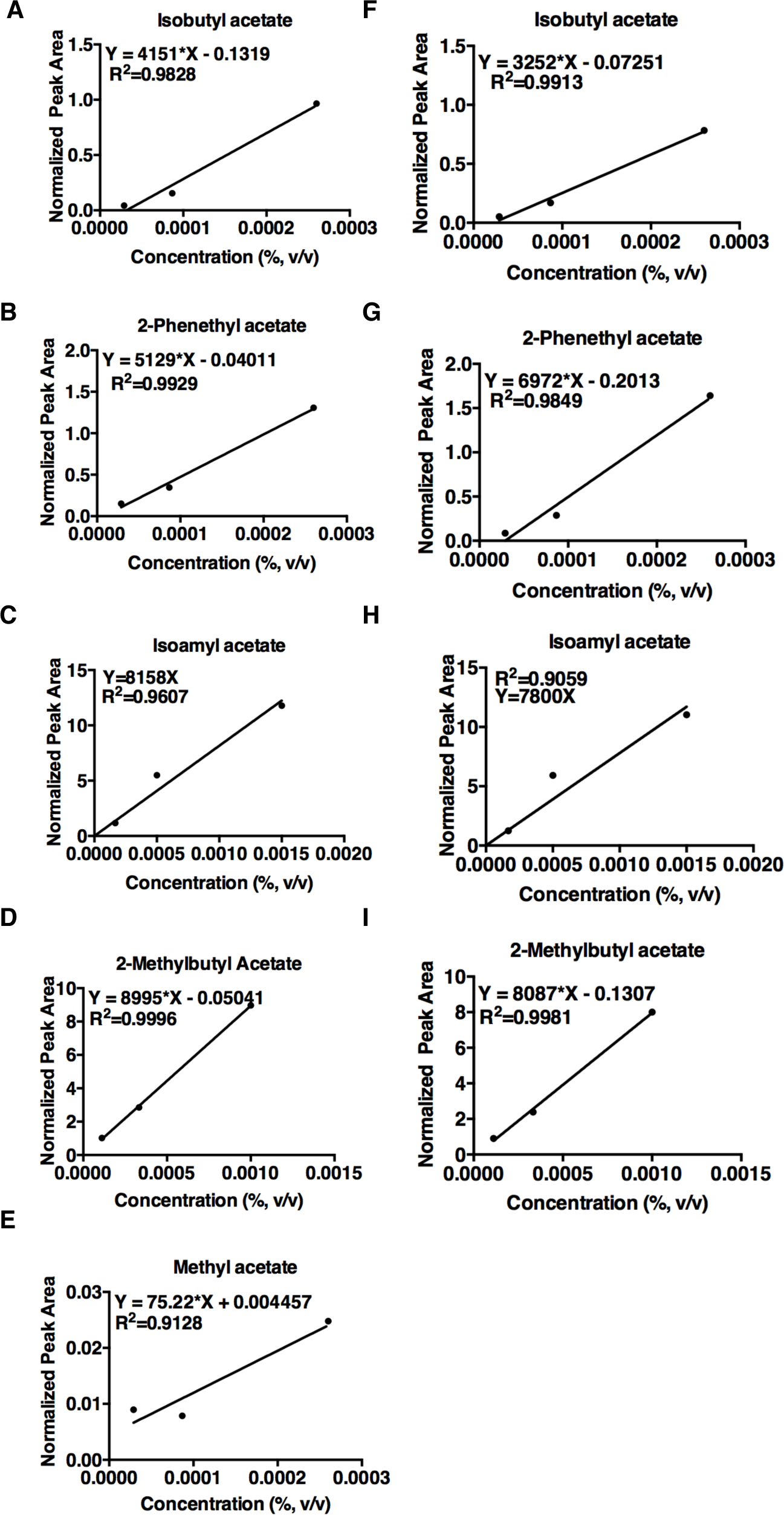
Normalized peak areas corresponding to metabolites in a defined metabolite mixture (from SPME GC-MS). A linear regression was calculated to quantify the metabolites in the co-culture. Each concentration is from one replicate. **A-E** and **F-I** are two separate experiments. Linear regression was used to estimate the concentration of the metabolites in the co-culture containing *S. cerevisiae* and *A. malorum* (Table 2) and to complement the co-culture containing *A. pomorum adhA* (Figure 5A).

**Figure 6-figure supplement 1.**
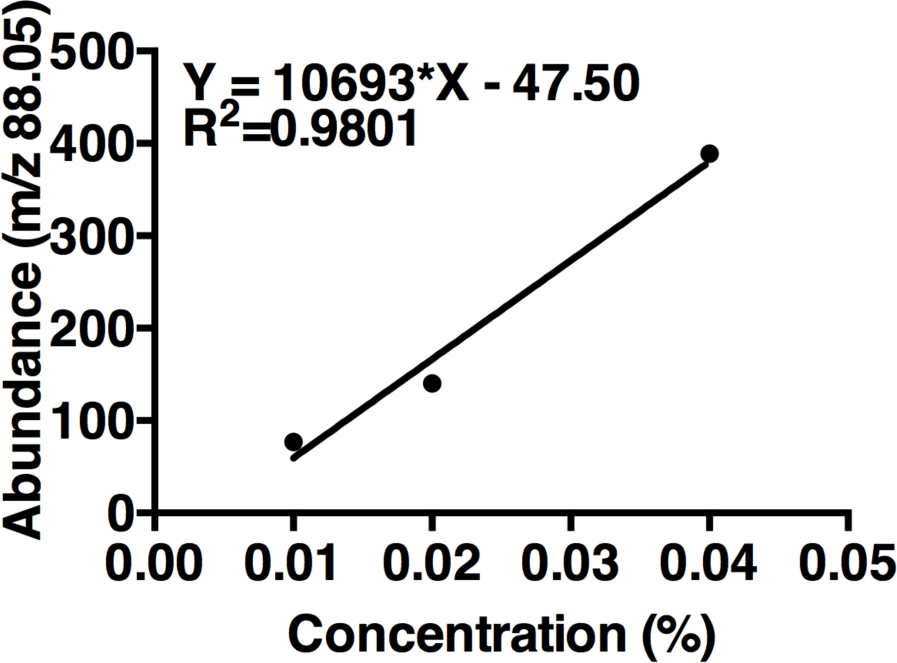
Acetoin linear regression. The curve is based on maximum m/z values (88.05) of three concentrations of acetoin. One replicate per concentration (n=1 experiment). The linear regression was used to estimate acetoin concentrations in the tri-culture and the co-culture (Figure 6B).

**Figure 6-figure supplement 2.**
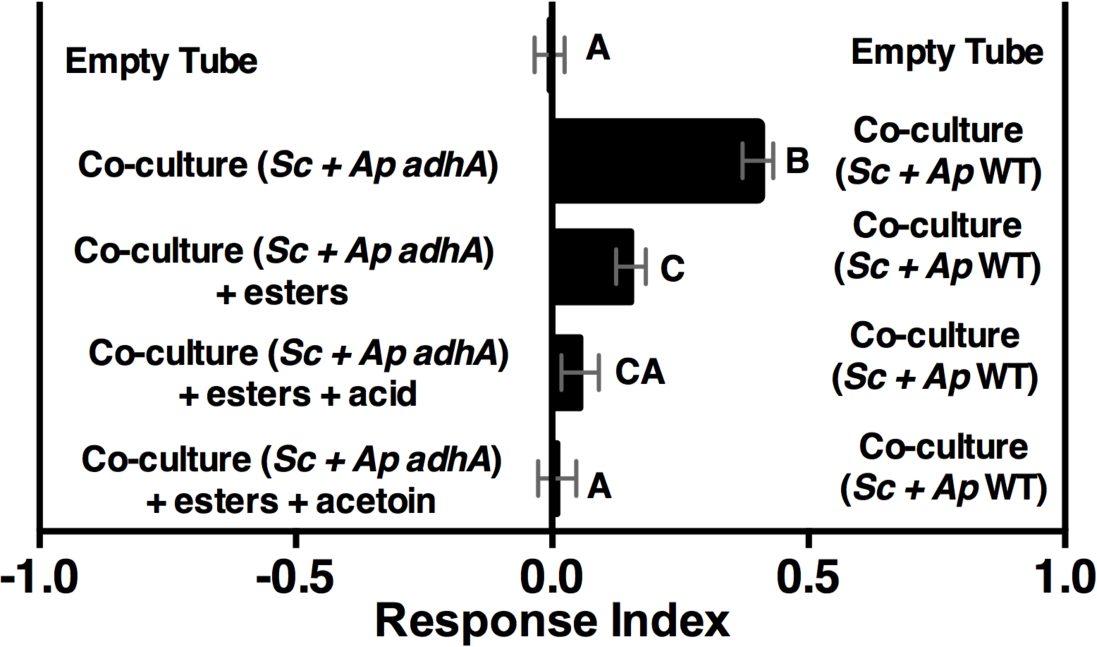
Role of esters alone or esters with acetoin in complementing co-culture with *A. pomomrum adhA*. Mean +/- SEM of three experiments with 16-18 total replicates. A one-way ANOVA assessed whether groups were different. A Tukey’s multiple comparison test was used to assess significant differences between groups (p≤0.05), as indicated by unique letters. Esters include ethyl acetate, isoamyl acetate, 2-phenethyl acetate, isobutyl acetate, 2-methylbutyl acetate, and methyl acetate. Esters added are based on physiological amounts in co-cultures and are calculated in Table 2 and Table 2-figure supplement 2). Acetoin is added in a similar amount as the tri-culture (Figure 6A, 0.3%). *Sc = S. cerevisiae, Ap = A. pomorum*.

**Figure 6-figure supplement 3.**
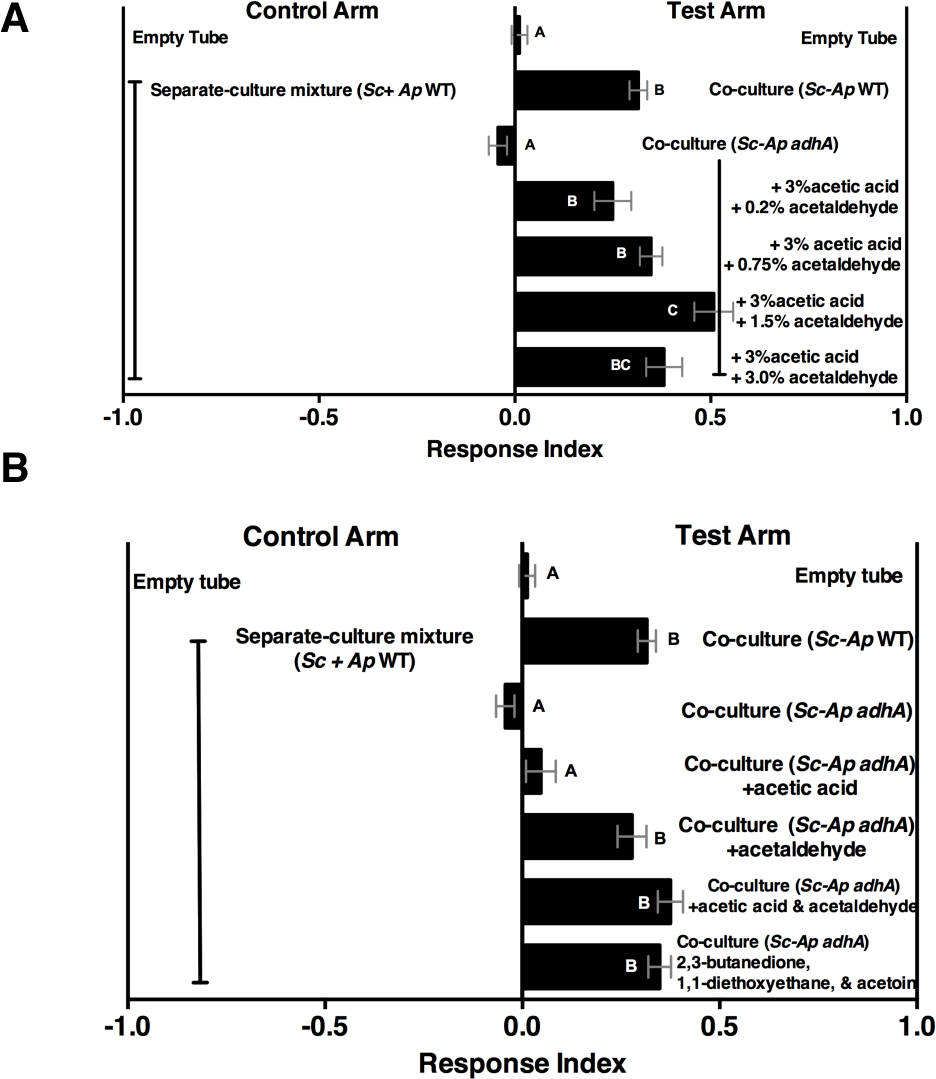
Acetaldehyde metabolic derivatives can complement the co-culture containing *A. pomorum adhA*, although their physiological concentrations are unknown. **(A)** Dose response of acetaldehyde was given to the co-culture containing *A. pomorum adhA* (along with a constant dose of 3.0% acetic acid). Metabolite additions were added to the culture in the noted percentages, allowed to sit 35 minutes at room temperature, mixed 1:1 in water, and assessed for *Drosophila* attractivess. Mean ± SEM of 1-7 two separate groups during the day. Differences between groups were tested by one-way ANOVA. A Tukey’s experiments with 12-42 total replicates per group. The experimental groups from one experiment were run in two separate groups during the day. Differences between groups were tested by one-way ANOVA. A Tukey’s multiple comparison test was used to assess significant differences between groups (p≤0.05), as indicated by unique letters. **(B)** Role of acetic acid, acetaldehyde, or specified acetaldehyde metabolic derivatives in complementing the co-culture containing *A. pomorum adhA.* Acetic acid (3.0%) and acetaldehyde (0.75%) were added and allowed to sit at room temperature for 35 minutes, mixed 1:1 in water, and *Drosophila* attraction was assayed. 2,3-butanedione (0.15%), 1,1-diethoxyethane (0.01%), and acetoin (0.15%) were added to the culture, mixed 1:1 with water, and *Drosophila* behavior was assayed. Mean ± SEM of 2-7 experiments with 5-6 replicates per experiment. Differences between groups were tested by one-way ANOVA. A Tukey’s multiple comparison test was used to assess significant differences between groups (p≤0.05), as indicated by unique letters. *Sc = S. cerevisiae, Ap = A. pomorum*.

**Figure 6-figure supplement 4.**
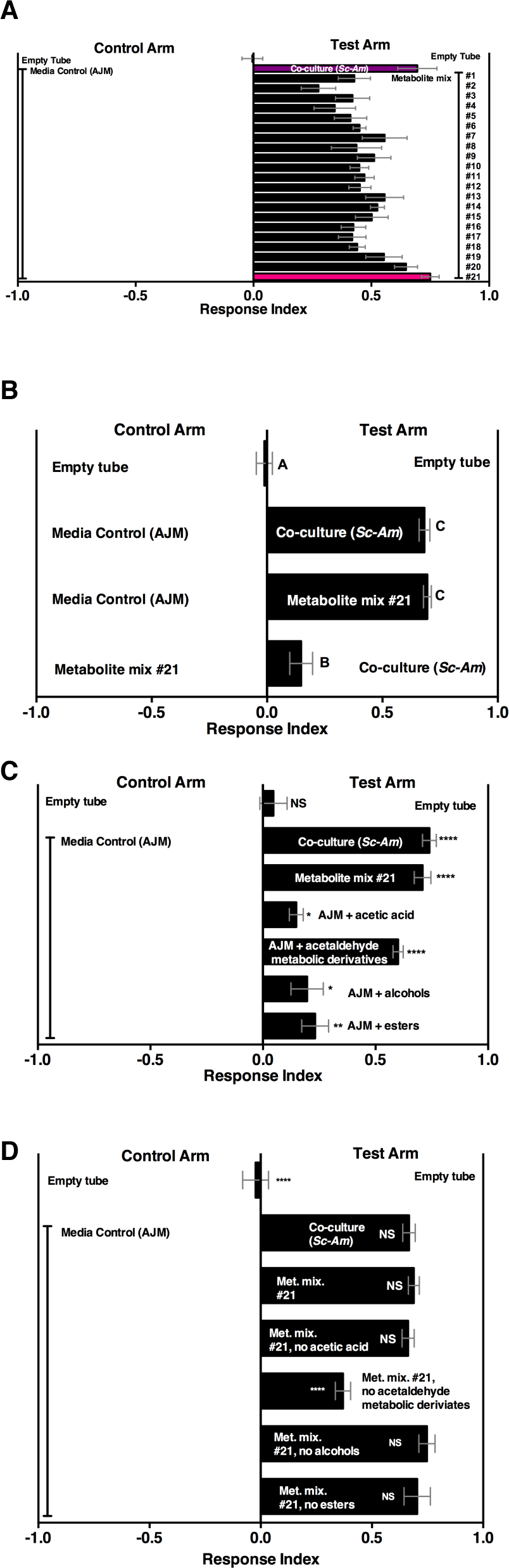
*Drosophila* behavior toward metabolite mixtures and their component parts **(A)** *Drosophila* behavior toward 21 metabolite mixtures versus AJM (50% in water). Methods Table 4 contains the concentrations of all mixtures (in 50% AJM). The co-culture (*S. cerevisiae* and *A. malorum*, purple bar) was grown for 96 h. The pink bar is metabolite mixture #21 and most closely resembled the co-culture. Mean ± SEM of 4-6 replicates per experimental group. Groups were run over five different days. **(B)** Relative *Drosophila* attraction to a co-culture grown for 96 h and metabolite mixture #21. Mean ± SEM of three experiments with 17-18 replicates per group. One-way ANOVA assessed whether the groups were different. A Tukey’s multiple comparison test was used to assess significant differences between groups (p≤0.05), as indicated by unique letters. **(C)** The sufficiency of individual groupings within metabolite mixture #21 to attract *Drosophila*. The individual grouping are as follows: acetaldaldehyde metabolic derivatives (1,1-diethoxyethane; acetoin; 2,3-butanedione); alcohols (ethanol; isobutanol; isoamyl alcohol; 2-methyl, 1-butanol; benzeneethanol); esters (isoamyl acetate; ethyl acetate; isobutyl acetate; 2-phenethyl acetate; butyl acetate; 2-methylbutyl acetate; methyl acetate; phenethyl benzoate; propyl acetate; ethyl isobutyrate; ethyl hexanoate; isovaleric acid; butyl ester; ethyl octanoate; ethyl decanoate; ethyl laurate); and acetic acid (acetic acid). Concentrations are in Methods Table 4. Mean ± SEM of six replicates of one experiment (except the acetaldehyde metabolic derivative group which is 12 replicates from 2 experiments). A t-test against a hypothetical value of 0 was performed. p >0.05; * p ≤0.05; ** p ≤0.01; *** p ≤0.001; **** p ≤0.0001 **(D)** The same groups used in **C** were used and removed from metabolite mix #21. The difference between the co-culture (*Sc-Am*) and each group was assessed by oneway ANOVA. A Tukey’s multiple comparison test was used to assess significant differences between groups (p≤0.05), as indicated by unique letters. Mean ± SEM of six replicates of one experiment are shown.

**Figure 6-figure supplement 5.**
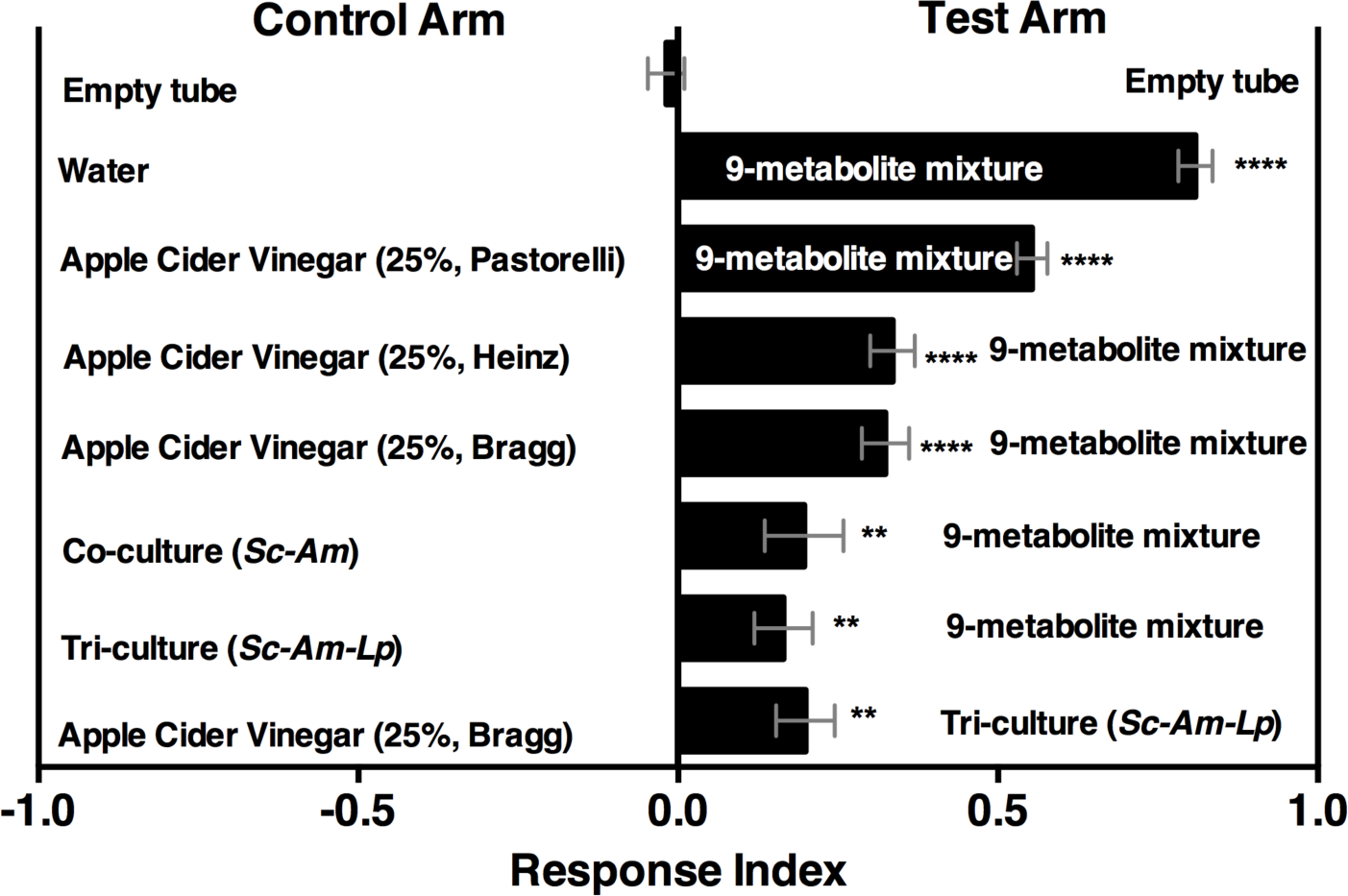
*Drosophila* behavior toward water amended with 9 metabolites (9-metabolite mixture) versus 3 different apple cider vinegars (ACV), a co-culture (*Sc-Am = S. cerevisiae* and *A. malorum*), or tri-culture (*Sc-Am-Lp = S. cerevisiae, A. malorum*, *L. plantarum* cs). Cultures were grown for 72 h and mixed 1:1 with water, as in all other experiments. Data points represent the Mean ± SEM of two experiments with twelve total replicates. A one-way ANOVA assessed whether each group was different from the mock condition (empty tube versus empty tube). p >0.05; * p ≤0.05; ** p ≤0.01; *** p ≤0.001; **** p ≤0.0001. The acetoin concentration was similar to that calculated from the tri-culture (Figure 6A, 0.3%). The concentrations of all 9 metabolites can be found in the materials and methods.

## MATERIALS AND METHODS

### Fly maintenance

Fly stocks, genotypes, and sources are listed in Methods Table 1. *Drosophila melanogaster* was reared at 25°C on a 12h:12h light: dark cycle on autoclaved food (5% yeast, 10% dextrose, 7% cornmeal, 0.6% propionic acid, 0.7% agar).

### Microbial strains

Microorganisms used in this study are listed and described in table S1. Microorganisms were streaked onto yeast-peptone dextrose (YPD; 1% yeast extract (Becton Dickinson, and Company, Franklin Lakes, NJ, USA), 2% peptone (Becton Dickinson, and Company, Franklin Lakes, NJ, USA), and 2% dextrose (Avantor Performance Materials, Center Valley, PA, USA)) or Man, de Rosa, Sharpe (MRS, Fisher Scientific, Waltham, MA, USA) plates from a freezer stock.

### T-maze olfactory attraction assays

The T-maze apparatus was a kind gift of the Carlson Laboratory. Flies were wet-starved for 15-26 hours prior to T-maze olfactory experiments by placing flies into vials containing Kimwipes (Kimberly Clark, Dallas, TX, USA) soaked with 2 mL of milliQ water. Flies were collected within 4 days (<65 flies per vial) of emergence and matured on autoclaved food. Flies between 3 and 10 days-old were used in experiments.

Single microbial colonies were picked from rich media (MRS and YPD) plates and grown overnight. Cultures were washed 1X in PBS, diluted 100-fold, and 10µl was aliquoted into 3mL of apple juice media (AJM, apple juice (Martinelli’s Gold Medal, Watsonville, CA, USA), pH adjusted to 5.3 with 5M NaOH, with 0.5% yeast extract). Media was filtered with a 0.22 µM-size pore attached to a 250-mL polystyrene bottle (Corning, NY, USA). For co-culture experiments, 1e3-1e5 CFU of each microorganism was placed simultaneously into AJM. Microorganisms were grown in 14-mL round bottom polypropylene tubes (Corning Science, Tamaulipas, Mexico) at 28°C, 200rpm for the time noted in individual experiments. The microbial culture was diluted 1:1 with sterile milliQ water (0.22 µM filter (Millipore, Billerica, MA, USA)) and placed directly onto autoclaved 10 mM round Whatman filter paper (GE Healthcare Life Science, Pittsburgh, PA, USA) placed near the bottom of 15mL CentriStar centrifuge tubes (Corning, NY). A total volume of 10 µl was used for all experiments.

Tubes containing microorganisms and *Drosophila* were placed into the behavioral room (20-25°C, 50-70% humidity maintained by a humidifier (Sunbeam Tower Humidifier, Boca Raton, FL, USA) and equilibrated for 10 minutes prior to the beginning of the experiment. Flies (∼40-130) were knocked into the T-maze apparatus, rested for ∼1 min, and then allowed to choose from the test and control arms for 2 min in the dark. The test arm was alternated from one side of the apparatus to the other every experimental replicate. A Response Index (RI) was computed to analyze preference for the test arm (flies in test arm-flies in control arm)/(total flies).

### Chemicals

Chemicals can be found in Methods table 2.

### Microbial populations and pH

Selective plates were used to distinguish *S. cerevisiae* from *A. malorum*. MRS containing 50 µg/mL cycloheximide selected for *A. malorum* while MRS containing 10 µg/mL chloramphenicol and 20 µg/mL tetracycline selected for *S. cerevisiae*. pH of filtered cultures (0.22 µM) was measured using a Beckman Coulter pH meter (Model Phi510, Fullerton, CA, USA).

### Gas chromatography-Mass spectrometry

Microbial samples were grown in AJM for a specified amount of time in 14-mL round bottom tubes fitted with an autoclaved tissue strainer (250 µM nylon mesh (Thermo Scientific Pierce, Grand Island, NY) holding between 0.03 and 0.05 grams of autoclaved Amberlite XAD-4 resin (Sigma-Aldrich, St Louis, MO, USA) prewashed in water and methanol. After microbial growth, XAD-4 from two cultures was dumped into an autoclaved glass vial. XAD-4 was swirled with 900µl methanol for 30 seconds. 500-750µl of methanol was removed for GC-MS analysis. Quantification for table S2 was derived from beads suspended above the cultures from 84-96 h of growth. Quantification for Fig. 2D was derived from beads suspended above the culture every 12 h; time points on the graph refer to the end point of the 12-h span (e.g. 84 h corresponds to beads suspended from 72-84 h of growth).

Samples (5 µl of methanol-extracted samples) were injected into the GC-MS (Agilent 7890A/5975C) at 250 °C using helium as the carrier gas at a flow rate of 1.1 mL per minute (column head pressure 13 psi). The following chromatography temperature program was used for experiments to initially identify metabolites in the co-culture and individually grown microorganisms: 40°C for 3 min ramped at 1.7°C per minute to 200 °C (held for 3 min) then to 220°C at 3 °C per min and held for a further 5 min. The total run time was 111.78 min. For experiments focused on the five major metabolites, a shorter program was used that maintained the same first 10 min of the previous method (all 5 volatiles eluted within 9 min). The chromatography temperature program was 40°C for 3 min, ramped at 1.7°C per min to 46.8°C and held for 3 min, then ramped at 60°C per min until 220°C and held for 5 min. The total run time was 17.9 min.

The mass spectrometer was run in electron-impact (EI) mode at 70 eV. The temperatures of the transfer line, quadrupole, and ionization source were 220°C, 180°C, and 230°C respectively. The ionization was off during the first 4 min to avoid solvent overloading with a source temperature of 230°C. Mass spectra were recorded in the range of 35-300 m/z. Full scan mode was used at a scan rate of 6 scans/sec. The electron multiplier voltage was set in the relative mode to autotune procedure.

In the initial experiments peaks were manually picked using Agilent Chemstation Software. Volatiles associated with peaks were searched against the National Institute of Standards (NIST) 11 database. Subsequent experiments focused on the 5 major volatiles identified in the initial experiments by performing extracted ion chromatograms using an ion that successfully identified a standard at a specific retention time. Quantification was performed by tabulating the maximum abundance of the ion at a characteristic retention time and using a linear regression equation from a dose-response of the standards (fig. S6).

### Headspace solid phase microextraction (SPME) Gas chromatography-Mass spectrometry

A Waters GCT Premier gas chromatography time of flight mass spectrometer (Milford, MA) with a DB-5MS column (30m x 0.25 mm ID x 0.25 μm film thickness; Agilent) was used. Live cultures were transferred to autoclaved glass vials (20 mL, 23 X 75 mm, Supelco, Bellefonte, PA, USA) with screw caps (18mm, 35 Shore A, Supelco, Bellefonte, PA, USA) after growing for 72 h.

The glass vials containing live microbial cultures were analyzed via a 50/30μm carboxen/divinylbenzene/polydimethylsiloxane Stableflex solid-phase micro-extraction (SPME) fiber. The extraction methodology was based on previous studies using SPME to extract volatiles form vinegars (65, 66). The syringe was inserted through the membrane of the caps and sampled the volatiles for 30 min at 45°C; subsequently, metabolites were desorbed for 30 sec at 240C and baked for an additional 4.5 min in the injection port. The gas chromatograph was fitted with a microchannel plate (MCP) detector. The temperature program of the column was as follows: 40°C for 5 min, 2°C a min for 17.5 min followed by 25°C a min for 10 min. A final hold time of 5 min at 325°C was used. The carrier gas was helium. A split ratio of 250 was used based on better peak resolution. An internal standard of cineole (Sigma-Aldrich, St. Louis, MO, USA) was run with each sample and used to compute relative abundances. The mass detector was in the range of 40 to 650 m/z.

To analyze the data, MassLynx software was used. The response threshold was set to an absolute area of 10.00. The software automatically picked out peaks and computed peak areas. To obtain a relative quantification, peaks were compared across samples and normalized to the internal standard. Peaks were first searched against the NIST5 database to identify potential hits. Most potential metabolites were confirmed by a standard mixture in 50% AJM. The standard mixtures are in Methods Table 3.

### Chemical complementation of co-culture containing A. pomorum PQQ-ADH-I

A co-culture containing *S. cerevisiae* and *A. pomorum* WT or *A. pomorum PQQ-ADH-I* was grown for 72 h before use in the T-maze. For the physiological concentrations of acetate-derived metabolites, concentrations were added as in table S3 and then mixed 1:1 with water prior to behavioral analysis. For the acetaldehyde metabolic derivatives chemical complementation group, a 1:1 mixture of the mutant co-culture: water was supplemented with 1,1-diethoxyethane, 2,3-butanedione, and acetoin at final concentrations of 0.01%, 0.15% and 0.15%, respectively and used immediately in the T-maze. Acetic acid and/or acetaldehyde were added to the culture, allowed to sit at RT for 35 min, mixed 1:1 with water and then placed into the T-maze vials.

Standard curves were used to calculate the concentrations of individual metabolites (fig. S10). The standard curves were generated on 2 separate experiments in which 3 concentrations of each standard was used. The concentrations of the metabolites were independently calculated from the standard curve equations generated on the two separate days. Estimated concentrations from each standard curve equation were averaged (table S3). The experimental data is based on the peak areas of the *S.cerevisiae-A. malorum* co-culture.

### Ester, acid, and acetaldehyde metabolic derivative mixture

The 9-metabolite mixture contains 1.5% acetic acid, 0.0003% isoamyl acetate, 0.0003% 2-phenethyl acetate, 0.01% ethyl acetate, 0.002% ethyl lactate, 0.3% 1,1-diethoxyethane, 0.3% 2,3-butanediol, 0.3% 2,3-butanedione, and 0.3% acetoin in filtered milliQ water.

### Drosophila survival in the presence of ethanol and acetic acid

Adult male flies (0-3 d-old) were collected and matured for 1 day on fly food. Flies were then placed into vials containing kimwipes with 5 mL of either Shields & Sang Insect Medium (Sigma, St. Louis, MO; positive control), MilliQ water (negative control), or MilliQ water with ethanol (9.4%), acetic acid (3.42%), or ethanol and acetic acid (1.4 and 2.8% respectively). Survival was assessed every 12 h for 7 d. For each condition 5 mL was given at 0 and 12 h and every 24 h thereafter. Experimental replicates were considered separate vials (5-6 per group). Each replicate contained 8-31 flies.

### Egg-laying preference assay

Egg-preference assay was adapted from Joseph *et al* 2009 (67). Microbial cultures grown for 96 h were heated to 65°C for 10 min, mixed 1:1 with 1.6% agarose and poured into a 35 mm petri dish separated in two by a straight-edge razor blade. Flies were starved for ∼18 h prior to the experiment. The 35 mm petri dish was placed within clear flat top boxes with dimensions 2 5/16” X 2 5/16” X 5 1/16” (TAP plastics, San Leandro, CA, USA). The test and control sides were alternated for each replicate. *Drosophila* aged 4-10 days (n=50-100) was allowed to lay eggs for 8 h. After the assay, the number of eggs on deposited on each choice was tabulated and an egg-laying index was computed analogously to the olfactory response index.

### Data analysis

Data analysis was performed in Prismv6.0b. Specific statistical tests are noted for individual experiments.

**Methods Table 1.** *Drosophila melanogaster* stocks used in experiments

| Notation | Gene<br>Mutant | Genotype | Source |
| --- | --- | --- | --- |
| CS | NA | w+-Canton-S | John Carlson Laboratory |
| Or83b | *Or83b* | w-Canton-S;;orco2 | John Carlson Laboratory |
| Ir64a | *Ir64a* | w[1118]; Mi{ET1}Ir64a[MB05283] | Bloomington Stock Center |
| Ir75a | *Ir75a* | y[1] w[67c23]; Mi{ET1}Ir75a[MB00253] | Bloomington Stock Center |
| W-CS | NA | w-Canton-S | John Carlson Laboratory |
| Balancer | NA | w-CS;Sp/Cyo;Dr/TM3B | John Carlson Laboratory |
| Ir25a | *Ir25a* | w<sup>1118</sup>; Mi{ET1}Ir25a<sup>MB09568</sup> | Bloomington Stock Center |
| Ir8a | *Ir8a* | Ir8a<sup>1</sup>; Bl<sup>1</sup> L<sup>2</sup>/CyO | Bloomington Stock Center |
| Ir76b | *Ir76b* | y1 w67c23; Mi{ET1}Ir76bMB00216 | Bloomington Stock Center |
| Or42b | *Or42b* | w; or42b/or42b; +/+ | John Carlson Laboratory |
| Or35a | *Or35a* | W; or35a/or35a; +/+ | John Carlson Laboratory |

| Chemical | Company | Location |
| --- | --- | --- |
| Paraffin Oil | EMD chemicals | Gibbstown, NJ, USA |
| Apple Cider Vinegar | Heinz | Pittsburgh, PA, USA |
| Apple Cider Vinegar<br>(unpasteurized) | Bragg (Fig. 3C only) | Santa Barbara, CA, USA |
| Apple Cider Vinegar | Pastorelli (Fig. 3C only) | Chicago, IL, USA |
| Benzaldehyde | Sigma Aldrich | St. Louis, MO, USA |
| 2-Methyl-1-Propanol (Isobutanol) | Sigma Aldrich | St. Louis, MO, USA |
| Isopentyl Acetate (Isoamyl Acetate) | Sigma Aldrich | St. Louis, MO, USA |
| Isoamyl Alcohol | Avantor Performance Materials | Center Valley, PA, USA |
| Acetic Acid, glacial | Avantor Performance Materials | Center Valley, PA, USA |
| Methanol | Avantor Performance Materials | Center Valley, PA, USA |
| Ethanol, 200 proof | Decon Laboratories | King of Prussia, PA, USA |
| Acetal (1,1, diethoxyethane) | Fisher | Pittsburgh, PA, USA |
| Acetaldehyde | Acros Organics | Grand Island, NY |
| Ethyl acetate | Sigma Aldrich | St. Louis, MO, USA |
| 1-pentanol | Sigma Aldrich | St. Louis, MO, USA |
| Benzeneethanol (2-phenylethanol) | Fisher | Pittsburgh, PA, USA |
| Isobutyl acetate | Sigma Aldrich | St. Louis, MO, USA |
| 2-phenethyl acetate | Sigma Aldrich | St. Louis, MO, USA |
| Butyl acetate | Sigma Aldrich | St. Louis, MO, USA |
| 2-methylbutyl acetate | TCI | Tokyo, Japan |
| Methyl acetate | Sigma Aldrich | St. Louis, MO, USA |
| Acetoin acetate | Sigma Aldrich | St. Louis, MO, USA |
| Phenethyl formate | Sigma Aldrich | St. Louis, MO, USA |
| Phenethyl benzoate | Sigma Aldrich | St. Louis, MO, USA |
| acetoin | Sigma Aldrich | St. Louis, MO, USA |
| 2,3-butanedione | Sigma Aldrich | St. Louis, MO, USA |
| 2-methyl-1-butanol | Sigma Aldrich | St. Louis, MO, USA |
| Ethyl lactate | Sigma Aldrich | St. Louis, MO, USA |
| 1-hexanol | Sigma Aldrich | St. Louis, MO, USA |
| Phenylacetic acid | Sigma Aldrich | St. Louis, MO, USA |
| 2,3-butanediol | Sigma Aldrich | St. Louis, MO, USA |
| Ethyl octanoate | Sigma Aldrich | St. Louis, MO, USA |
| Ethyl decanoate | Sigma Aldrich | St. Louis, MO, USA |
| Ethyl hexanoate | Sigma Aldrich | St. Louis, MO, USA |
| Ethyl laurate | Sigma Aldrich | St. Louis, MO, USA |
| Ethyl isobutyrate | Sigma Aldrich | St. Louis, MO, USA |
| phenylacetaldehyde | Sigma Aldrich | St. Louis, MO, USA |
| Butyl isovalerate | TCI | Tokyo, Japan |
| Propyl acetate | Sigma Aldrich | St. Louis, MO, USA |

**Methods Table 3.**
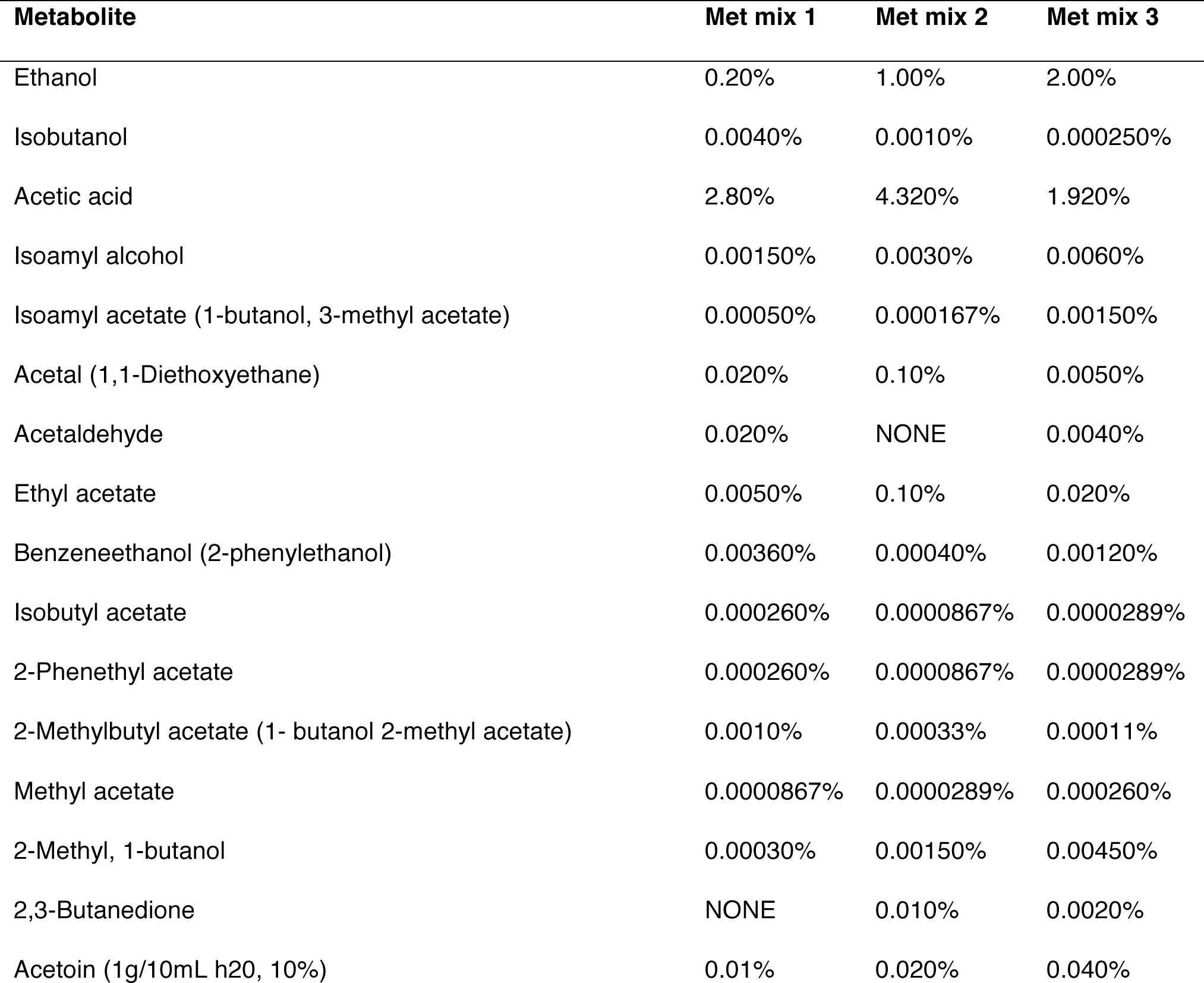
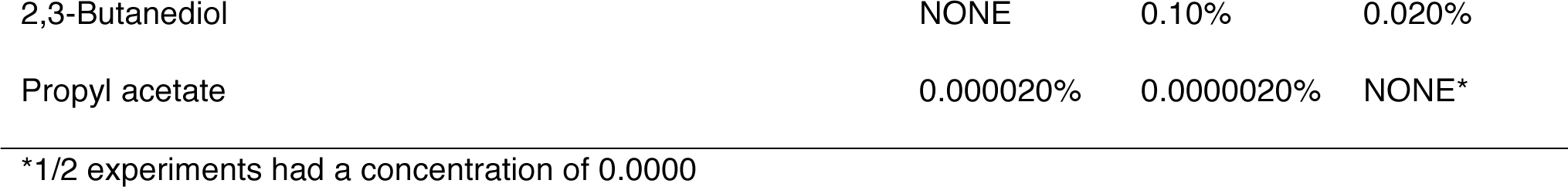
Metabolite mixture concentrations used for identification and quantification in SPME GC-MS

| Metabolite | Met mix 1 | Met mix 2 | Met mix 3 |
| --- | --- | --- | --- |
| Ethanol | 0.20% | 1.00% | 2.00% |
| Isobutanol | 0.0040% | 0.0010% | 0.000250% |
| Acetic acid | 2.80% | 4.320% | 1.920% |
| Isoamyl alcohol | 0.00150% | 0.0030% | 0.0060% |
| Isoamyl acetate (1-butanol, 3-methyl acetate) | 0.00050% | 0.000167% | 0.00150% |
| Acetal (1,1-Diethoxyethane) | 0.020% | 0.10% | 0.0050% |
| Acetaldehyde | 0.020% | NONE | 0.0040% |
| Ethyl acetate | 0.0050% | 0.10% | 0.020% |
| Benzeneethanol (2-phenylethanol) | 0.00360% | 0.00040% | 0.00120% |
| Isobutyl acetate | 0.000260% | 0.0000867% | 0.0000289% |
| 2-Phenethyl acetate | 0.000260% | 0.0000867% | 0.0000289% |
| 2-Methylbutyl acetate (1- butanol 2-methyl acetate) | 0.0010% | 0.00033% | 0.00011% |
| Methyl acetate | 0.0000867% | 0.0000289% | 0.000260% |
| 2-Methyl, 1-butanol | 0.00030% | 0.00150% | 0.00450% |
| 2,3-Butanedione | NONE | 0.010% | 0.0020% |
| Acetoin (1g/10mL h20, 10%) | 0.01% | 0.020% | 0.040% |
| 2,3-Butanediol | NONE | 0.10% | 0.020% |
| Propyl acetate | 0.000020% | 0.0000020% | NONE* |

\*1/2 experiments had a concentration of 0.0000

**Methods Table 4.**
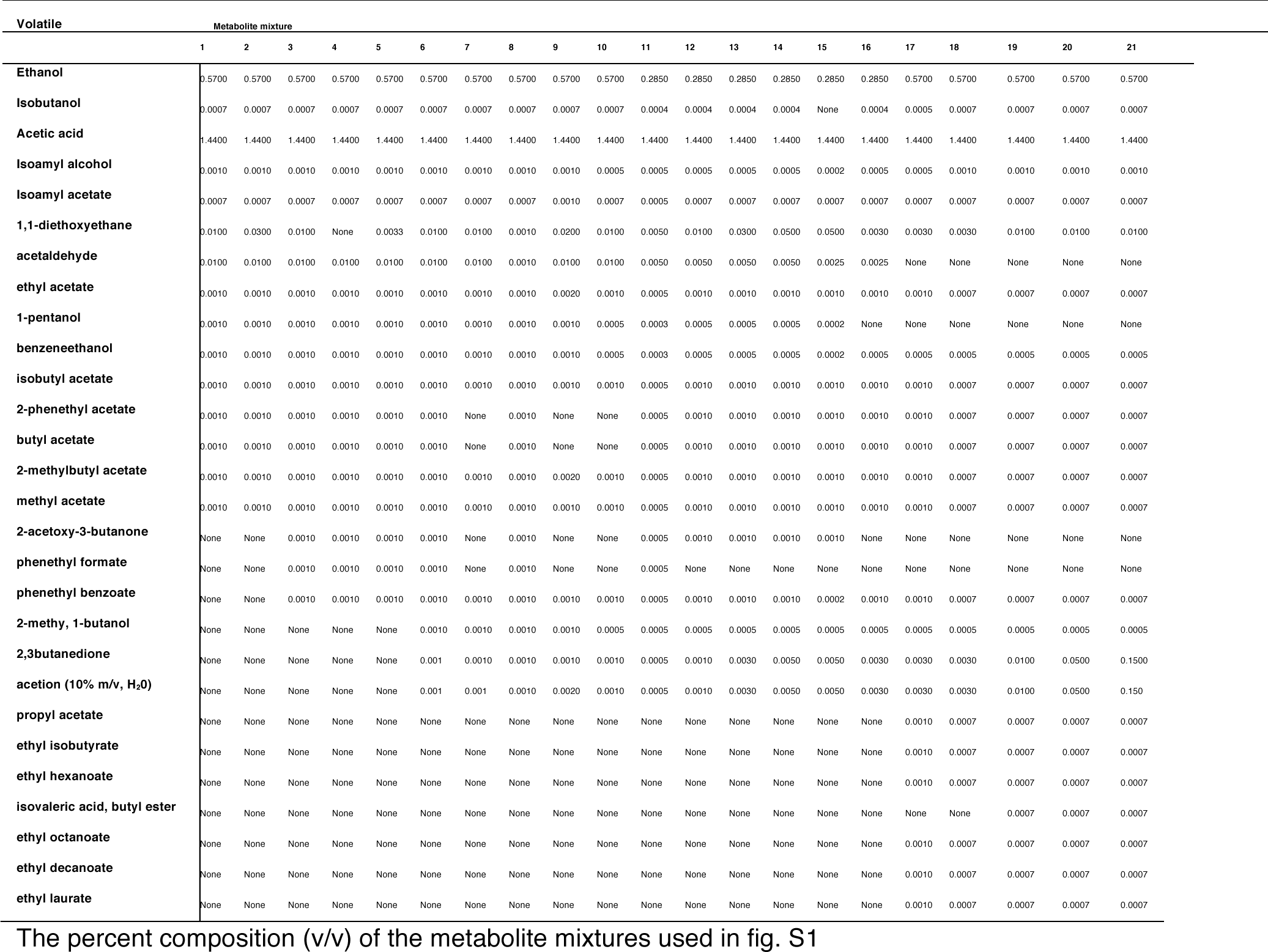
Composition of metabolite mixtures 1-21 used in figure S14

## References

1. Mcfall-Ngai M, Hadfield MG, Bosch TCG, Carey HV, Domazet-Loso T, Douglas AE, Dubilier N, Eberl G, Fukami T, Gilbert SF, Hentschel U, King N, Kjelleberg S, Knoll AH, Kremer N, Mazmanian SK, Metcalf JL, Nealson K, Pierce NE, Rawls JF, Reid A, Ruby EG, Rumpho M, Sanders JG, Tautz D, Wernegreen JJ. 2013. Animals in a bacterial world, a new imperative for the life sciences. PNAS 110:3229–3236.

2. van Nood E, Vrieze A, Nieuwdorp M, Fuentes S, Zoetendal EG, de Vos WM, Visser CE, Kuijper EJ, Bartelsman JFWM, Tijssen JGP, Speelman P, Dijkgraaf MGW, Keller JJ. 2013. Duodenal infusion of donor feces for recurrent *Clostridium difficile*. N Engl J Med 368:407–415.

3. Zhu L, Wu Q, Dai J, Zhang S, Wei F. 2011. Evidence of cellulose metabolism by the giant panda gut microbiome. PNAS 108:17714–17719.

4. Degnan PH, Taga ME, Goodman AL. 2014. Vitamin B12 as a modulator of gut microbial ecology. Cell Metab 20:769–778.

5. Bravo JA, Forsythe P, Chew MV, Escaravage E, Savignac HM, Dinan TG, Bienenstock J, Cryan JF. 2011. Ingestion of Lactobacillus strain regulates emotional behavior and central GABA receptor expression in a mouse via the vagus nerve. PNAS 108:16050–16055.

6. Rath CM, Dorrestein PC. 2012. The bacterial chemical repertoire mediates metabolic exchange within gut microbiomes. Curr Opin Microbiol 15:147–154.

7. Manor O, Levy R, Borenstein E. 2014. Mapping the inner workings of the microbiome: genomic- and metagenomic-based study of metabolism and metabolic interactions in the human microbiome. Cell Metab 20:742–752.

8. Gerber GK. 2014. The dynamic microbiome. FEBS Letters 588:4131–4139.

9. Gonzalez A, King A, Robeson MS, Song S, Shade A, Metcalf JL, Knight R. 2012. Characterizing microbial communities through space and time. Current Opinion in Biotechnology 23:431–436.

10. Derewacz DK, Covington BC, McLean JA, Bachmann BO. 2015. Mapping microbial response metabolomes for induced natural product discovery. ACS chemical biology 10:1998–2006.

11. Goh E-B, Yim G, Tsui W, McClure J, Surette MG, Davies J. 2002. Transcriptional modulation of bacterial gene expression by subinhibitory concentrations of antibiotics. PNAS 99:17025–17030.

12. Ridaura VK, Faith JJ, Rey FE, Cheng J, Duncan AE, Kau AL, Griffin NW, Lombard V, Henrissat B, Bain JR, Muehlbauer MJ, Ilkayeva O, Semenkovich CF, Funai K, Hayashi DK, Lyle BJ, Martini MC, Ursell LK, Clemente JC, Van Treuren W, Walters WA, Knight R, Newgard CB, Heath AC, Gordon JI. 2013. Gut microbiota from twins discordant for obesity modulate metabolism in mice. Science 341:1241214.

13. Costello EK, Lauber CL, Hamady M, Fierer N, Gordon JI, Knight R. 2009. Bacterial community variation in human body habitats across space and time. Science 326:1694–1697.

14. Ivanov II, Atarashi K, Manel N, Brodie EL, Shima T, Karaoz U, Wei D, Goldfarb KC, Santee CA, Lynch SV, Tanoue T, Imaoka A, Itoh K, Takeda K, Umesaki Y, Honda K, Littman DR. 2009. Induction of intestinal Th17 cells by segmented filamentous bacteria. Cell 139:485–498.

15. Poulsen M, Hu H, Li C, Chen Z, Xu L, Otani S, Nygaard S, Nobre T, Klaubauf S, Schindler PM, Hauser F, Pan H, Yang Z, Sonnenberg ASM, de Beer ZW, Zhang Y, Wingfield MJ, Grimmelikhuijzen CJP, de Vries RP, Korb J, Aanen DK, Wang J, Boomsma JJ, Zhang G. 2014. Complementary symbiont contributions to plant decomposition in a fungus-farming termite. PNAS 111:14500–14505.

16. Rakoff-Nahoum S, Coyne MJ, Comstock LE. 2014. An ecological network of polysaccharide utilization among human intestinal symbionts. Current Biology 24:40–49.

17. Rakoff-Nahoum S, Foster KR, Comstock LE. 2016. The evolution of cooperation within the gut microbiota. Nature 533:255–259.

18. Fraune S, Anton-Erxleben F, Augustin R, Franzenburg S, Knop M, Schröder K, Willoweit-Ohl D, Bosch TC. 2015. Bacteria-bacteria interactions within the microbiota of the ancestral metazoan *Hydra* contribute to fungal resistance. ISME J 9:1543–1556.

19. Chandler JA, Lang JM, Bhatnagar S, Eisen JA, Kopp A. 2011. Bacterial communities of diverse *Drosophila* species: ecological context of a host-microbe model system. PLoS Genet 7:e1002272.

20. Chandler JA, Eisen JA, Kopp A. 2012. Yeast communities of diverse *Drosophila* species: comparison of two symbiont groups in the same hosts. Applied and Environmental Microbiology 78:7327–7336.

21. Broderick NA, Lemaitre B. 2012. Gut-associated microbes of *Drosophila melanogaster*. Gut Microbes 3:307–321.

22. Camargo R, Phaff HJ. 1957. Yeasts occurring in *Drosophila* flies and in fermenting tomato fruits in northern California. J Food Science 22:367–372.

23. Barata A, Malfeito-Ferreira M, Loureiro V. 2012. The microbial ecology of wine grape berries. Int J Food Microbiol 153:243–259.

24. Erkosar B, Storelli G, Defaye A, Leulier F. 2013. Host-intestinal microbiota mutualism: “learning on the fly”. Cell Host & Microbe 13:8–14.

25. Blum JE, Fischer CN, Miles J, Handelsman J. 2013. Frequent replenishment sustains the beneficial microbiome of *Drosophila melanogaster*. MBio 4:e00860–13.

26. Broderick NA, Buchon N, Lemaitre B. 2014. Microbiota-induced changes in *Drosophila melanogaster* host gene expression and gut morphology. MBio 5:e01117–14.

27. Becher PG, Flick G, Rozpędowska E, Schmidt A, Hagman A, Lebreton S, Larsson MC, Hansson BS, Piškur J, Witzgall P, Bengtsson M. 2012. Yeast, not fruit volatiles mediate *Drosophila melanogaster* attraction, oviposition and development. Functional Ecology 26:822–828.

28. Christiaens JF, Franco LM, Cools TL, De Meester L, Michiels J, Wenseleers T, Hassan BA, Yaksi E, Verstrepen KJ. 2014. The fungal aroma gene *ATF1* promotes dispersal of yeast cells through insect vectors. Cell Rep 9:425–432.

29. Schiabor KM, Quan AS, Eisen MB. 2014. *Saccharomyces cerevisiae* mitochondria are required for optimal attractiveness to *Drosophila melanogaster*. PLoS ONE 9:e113899.

30. Palanca L, Gaskett AC, Günther CS, Newcomb RD, Goddard MR. 2013. Quantifying variation in the ability of yeasts to attract *Drosophila melanogaster*. PLoS ONE 8:e75332.

31. Venu I, Durisko Z, Xu J, Dukas R. 2014. Social attraction mediated by fruit flies' microbiome. J Exp Biol 217:1346–1352.

32. Ai M, Min S, Grosjean Y, Leblanc C, Bell R, Benton R, Suh GSB. 2010. Acid sensing by the *Drosophila* olfactory system. Nature 468:691–695.

33. Wong CNA, Ng P, Douglas AE. 2011. Low-diversity bacterial community in the gut of the fruitfly *Drosophila melanogaster*. Environ Microbiol 13:1889–1900.

34. Shin SC, Kim S-H, You H, Kim B, Kim AC, Lee K-A, Yoon J-H, Ryu J-H, Lee W-J. 2011. *Drosophila* microbiome modulates host developmental and metabolic homeostasis via insulin signaling. Science 334:670–674.

35. Storelli G, Defaye A, Erkosar B, Hols P, Royet J, Leulier F. 2011. *Lactobacillus plantarum promotes Drosophila* systemic growth by modulating hormonal signals through TOR-dependent nutrient sensing. Cell Metab 14:403–414.

36. Buchon N, Broderick NA, Chakrabarti S, Lemaitre B. 2009. Invasive and indigenous microbiota impact intestinal stem cell activity through multiple pathways in *Drosophila*. Genes Dev 23:23332344.

37. Newell PD, Douglas AE. 2014. Interspecies interactions determine the impact of the gut microbiota on nutrient allocation in *Drosophila melanogaster*. Applied and Environmental Microbiology 80:788–796.

38. Wong AC-N, Dobson AJ, Douglas AE. 2014. Gut microbiota dictates the metabolic response of *Drosophila* to diet. J Exp Biol 217:1894–1901.

39. El-Zoghbi M. 1994. Biochemical changes in some tropical fruits during ripening. Food Chem 49:33–37.

40. Abu-Goukh A-BA, Bashir HA. 2003. Changes in pectic enzymes and cellulase activity during guava fruit ripening. Food Chem 83:213–218.

41. Mao WW, Kinsella JE. 1981. Amylase activity in banana fruit: properties and changes in activity with ripening. J Food Science 46:1400–1403.

42. Barbe JC, De Revel G, Joyeux A, Bertrand A, Lonvaud-Funel A. 2001. Role of botrytized grape micro-organisms in SO2 binding phenomena. J Appl Microbiol 90:34–42.

43. Turner SL, Ray A. 2009. Modification of CO2 avoidance behaviour in *Drosophila* by inhibitory odorants. Nature 461:277–281.

44. Anagnostou C, Dorsch M, Rohlfs M. 2010. Influence of dietary yeasts on *Drosophila melanogaster* life-history traits. Entomologia Experimentalis et Applicata 136:1–11.

45. Abuin L, Bargeton B, Ulbrich MH, Isacoff EY, Kellenberger S, Benton R. 2011. Functional architecture of olfactory ionotropic glutamate receptors. Neuron 69:44–60.

46. Hallem EA, Carlson JR. 2006. Coding of odors by a receptor repertoire. Cell 125:143–160.

47. Larsson MC, Domingos AI, Jones WD, Chiappe ME, Amrein H, Vosshall LB. 2004. *Or83b* encodes a broadly expressed odorant receptor essential for *Drosophila* olfaction. Neuron 43:703–714.

48. Mathew D, Martelli C, Kelley-Swift E, Brusalis C, Gershow M, Samuel ADT, Emonet T, Carlson JR. 2013. Functional diversity among sensory receptors in a *Drosophila* olfactory circuit. PNAS 110:E2134–43.

49. Asahina K, Louis M, Piccinotti S, Vosshall LB. 2009. A circuit supporting concentration-invariant odor perception in *Drosophila*. Journal of Biology 8:1–19.

50. Stokl J, Strutz A, Dafni A, Svatos A, Doubsky J, Knaden M, Sachse S, Hansson BS, Stensmyr MC. 2010. A deceptive pollination system targeting Drosophilids through olfactory mimicry of yeast. Current Biology 20:1846–1852.

51. Semmelhack JL, Wang JW. 2009. Select *Drosophila* glomeruli mediate innate olfactory attraction and aversion. Nature 459:218–223.

52. Linda F Chuang EBC. 1968. Biosynthesis of diacetyl in bacteria and yeast. J Bacteriol 95:2083–2089.

53. Ruyters S, Mukherjee V, Verstrepen KJ, Thevelein JM, Willems KA, Lievens B. 2015. Assessing the potential of wild yeasts for bioethanol production. J Ind Microbiol Biotechnol 42:3948.

54. Ranganathan S, Davis DG, Hood RD. 1987. Developmental toxicity of ethanol in *Drosophila melanogaster*. Teratology 36:45–49.

55. Azanchi R, Kaun KR, Heberlein U. 2013. Competing dopamine neurons drive oviposition choice for ethanol in *Drosophila*. PNAS 110:21153–21158.

56. Kacsoh BZ, Lynch ZR, Mortimer NT, Schlenke TA. 2013. Fruit flies medicate offspring after seeing parasites. Science 339:947–950.

57. Lhocine N, Ribeiro PS, Buchon N, Wepf A, Wilson R, Tenev T, Lemaitre B, Gstaiger M, Meier P, Leulier F. 2008. PIMS modulates immune tolerance by negatively regulating *Drosophila* innate immune signaling. Cell Host & Microbe 4:147–158.

58. Ryu J-H, Kim S-H, Lee H-Y, Bai JY, Nam Y-D, Bae J-W, Lee DG, Shin SC, Ha E-M, Lee W-J. 2008. Innate immune homeostasis by the homeobox gene caudal and commensal-gut mutualism in *Drosophila*. Science 319:777–782.

59. Paredes JC, Welchman DP, Poidevin M, Lemaitre B. 2011. Negative regulation by amidase PGRPs shapes the *Drosophila* antibacterial response and protects the fly from innocuous infection. Immunity 35:770–779.

60. Costechareyre D, Capo F, Fabre A, Chaduli D, Kellenberger C, Roussel A, Charroux B, Royet J. 2016. Tissue-specific regulation of *Drosophila* NF-KB pathway activation by peptidoglycan recognition protein SC. J Innate Immun 8:67–80.

61. Broderick NA. 2016. Friend, foe or food? Recognition and the role of antimicrobial peptides in gut immunity and Drosophila-microbe interactions. Philos Trans R Soc Lond, B, Biol Sci 371:20150295.

62. Piper MDW, Blanc E, Leitão-Gonçalves R, Yang M, He X, Linford NJ, Hoddinott MP, Hopfen C, Soultoukis GA, Niemeyer C, Kerr F, Pletcher SD, Ribeiro C, Partridge L. 2014. A holidic medium for *Drosophila melanogaster*. Nat Methods 11:100–105.

63. Jarosz DF, Brown JCS, Walker GA, Datta MS, Ung WL, Lancaster AK, Rotem A, Chang A, Newby GA, Weitz DA, Bisson LF, Lindquist S. 2014. Cross-kingdom chemical communication drives a heritable, mutually beneficial prion-based transformation of metabolism. Cell 158:1083–1093.

64. Staubach F, Baines JF, Künzel S, Bik EM, Petrov DA. 2013. Host species and environmental effects on bacterial communities associated with *Drosophila* in the laboratory and in the natural environment. PLoS ONE 8:e70749.

65. Callejón RM, González AG, Troncoso AM, Morales ML. 2008. Optimization and validation of headspace sorptive extraction for the analysis of volatile compounds in wine vinegars. Journal of Chromatography A 1204:93–103.

66. Xiao Z, Dai S, Niu Y, Yu H, Zhu J, Tian H, Gu Y. 2011. Discrimination of chinese vinegars based on headspace solid-phase microextraction-gas chromatography mass spectrometry of volatile compounds and multivariate analysis. J Food Science 76:1125–1135.

67. Joseph RM, Devineni AV, King IFG, Heberlein U. 2009. Oviposition preference for and positional avoidance of acetic acid provide a model for competing behavioral drives in *Drosophila*. PNAS 106:11352–11357.

